# Optimized husbandry and targeted gene-editing for the cnidarian *Nematostella vectensis*

**DOI:** 10.1101/2023.04.14.536874

**Authors:** João E. Carvalho, Maxence Burtin, Olivier Detournay, Aldine R. Amiel, Eric Röttinger

## Abstract

Optimized laboratory conditions for research models are crucial for the success of scientific projects. This includes the control of the entire life cycle, access to all developmental stages and maintaining stable physiological conditions. Reducing the life cycle of a research model can also enhance the access to biological material and speed up genetic tool development. Thus, we optimized the rearing conditions for the sea anemone *Nematostella vectensis*, a cnidarian research model to study embryonic and post-metamorphic processes, such as regeneration.

We adopted a semi-automated aquaculture system for *N. vectensis* and developed a dietary protocol optimized for the different life stages. Thereby, we increased spawning efficiencies and post-spawning survival rates, and considerably reduced the overall life cycle down to two months. To further improve the obtention of CRISPR-Cas9 mutants, we optimized the design of sgRNAs leading to full KO animals in F0 polyps using a single sgRNA. Finally, we show that NHEJ-mediated transgene insertion is possible in *N. vectensis*. In sum our study provides additional resources for the scientific community that uses or will use *N. vectensis* as a research model.

**Summary statement:** Optimized life cycle, in combination with efficient gene-editing approaches facilitates the establishment of genetic tools in *N. vectensis*, an emerging model for environmental stress response, regeneration, and longevity.

## Introduction

Selecting an adequate research model to address original questions is fundamental for the success of any scientific project. It requires the weighting of several different factors and advantages, specific to the selected model, such as the capacity to maintain the model in captivity, the knowledge on how to reproduce it, manipulate it (*e.g.,* genetically) and the ease of accessing its different life stages. Classically used and well-established research models such as *Drosophila melanogaster*, *Caenorhabditis elegans*, zebrafish or mice, benefit from a large amount of accumulated knowledge, tools, a large and well-organized community of scientists, well-established laboratory strains and experimental methods (Bradford et al., 2022; Cook et al., 2016; Davis et al., 2022; Gramates et al., 2022).

Unconventional research models hold the potential to tackle novel questions and approaches for the study of complex biological processes difficult to address using established models (Cook et al., 2016). However, unconventional research models present some challenges regarding their establishment and use. They require a significant effort, at the community level, to set up standard methods for maintaining, raising, as well as propagating animals in the laboratory to experimentally address the specific scientific questions or complex biological traits of interest, such as regeneration and aging/longevity.

The implementation of such non-conventional research models is nowadays essential to gain insight into the mechanisms underlying these complex biological traits at the cellular, molecular, and genetic level (Amiel et al., 2021; Arboleda et al., 2018; Ivankovic et al., 2019; Kassmer et al., 2019; Layden et al., 2016; Lechable et al., 2020; Mehta and Singh, 2019; Röttinger, 2021; Srivastava, 2022). Methods for the development and improvement of the entire life cycle in stable and reproducible laboratory conditions have recently been described for several marine vertebrates (Roux et al., 2020) and invertebrates (Carvalho et al., 2017; Gordon et al., 2020; Henry et al., 2020; Holland and Li, 2021; Lechable et al., 2020; Murabe et al., 2021; Soto-Àngel et al., 2022).

One of the emerging non-conventional research models to address intriguing biological traits is the cnidarian *Nematostella vectensis* (Figure 1A). This sea anemone (Cnidaria, Anthozoa) has been initially developed for evolutionary developmental biology (EvoDevo) questions (Chourrout et al., 2006; Fritzenwanker et al., 2007; Kusserow et al., 2005; Lee et al., 2007; Wikramanayake et al., 2003) and emerged more recently as a promising model to study whole-body regeneration (Amiel et al., 2015; Amiel et al., 2019; Bossert et al., 2013; Johnston et al., 2021; Passamaneck and Martindale, 2012; Reitzel et al., 2007; Warner et al., 2018), environmental stress-response (Ambrosone et al., 2014; Baldassarre et al., 2022; Elran et al., 2014; Klein et al., 2021; Reitzel et al., 2008), and aging (reviewed in Amiel et al., 2021).

**Figure 1.**
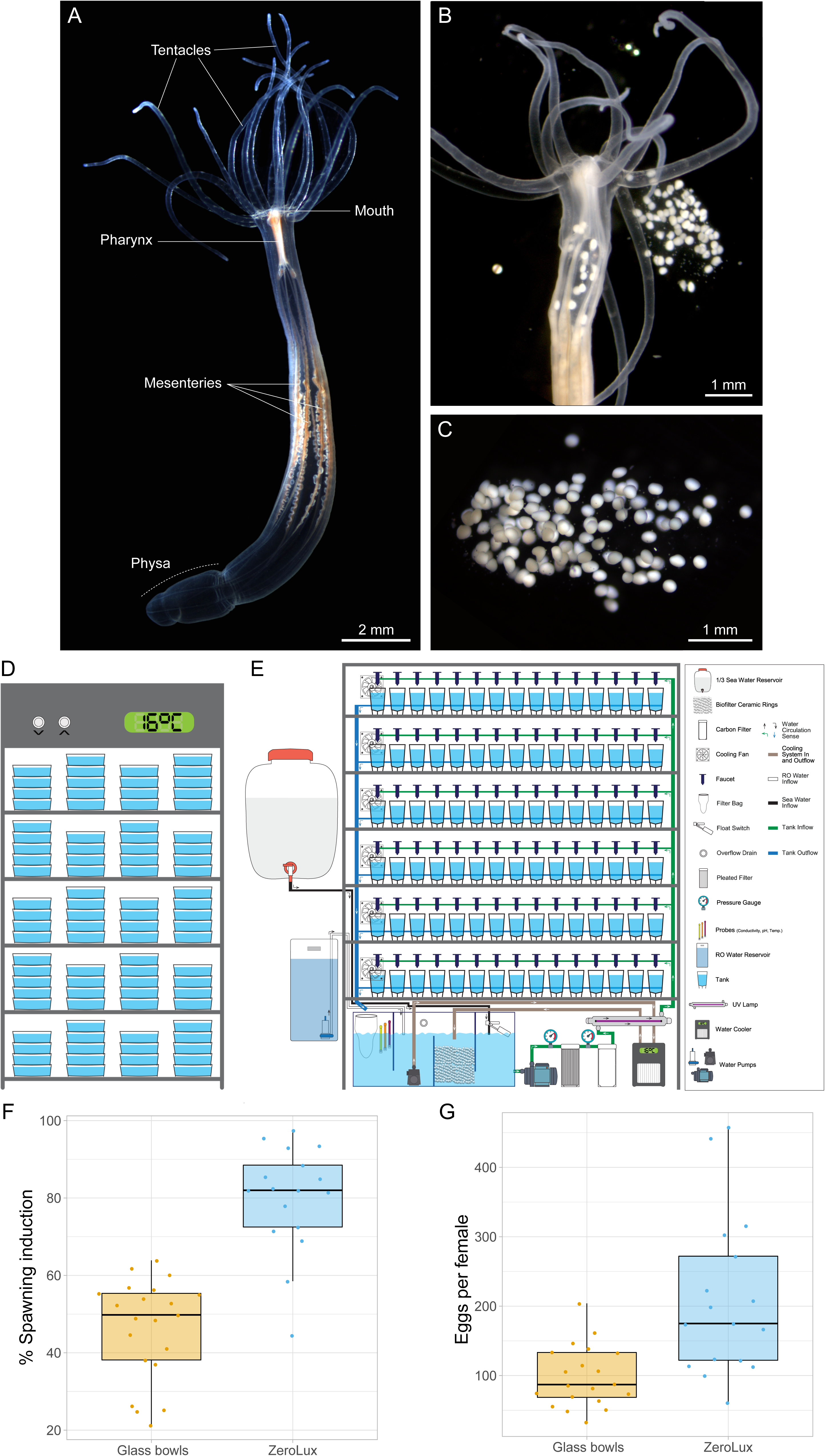
*Nematostella vectensis* spawning and optimized maintenance results. (A) Adult polyp of *N. vectensis*. (B) A spawning adult polyp of *N. vectensis* (C) Detailed view of an egg-mass. (D) Schematic overview of a *N. vectensis* maintenance incubator. (E) Schematic overview of *N. vectensis* of the semi-automated culture system, i.e. Zerolux. (F) Spawning induction percentage comparing *N. vectensis* maintenance in glass bowls and semi-automated aquaria system, over a period of 21 weeks. (G) Estimated mean number of eggs produced by *N. vectensis* females in glass bowls and Zerolux, over a period of 21 weeks. Welch’s t-test was used to verify the differences between both condition in D and E.

*N. vectensis* lives in estuarine environments like salt marshes or brackish water pools, it is gonochoric and able to reproduce sexually by external fertilization, as well as asexually via fission or physal pinching (Hand and Uhlinger, 1992). *N. vectensis* is rather simple to rear and when the polys are sexually mature, spawning and fertilization are controlled in laboratory condition leading to thousands of gametes and embryos every week (Figure 1B, C). Furthermore, the entire life-cycle can be closed in the laboratory, allowing for the development of large reproducible colonies from only a handful of polys (Fritzenwanker and Technau, 2002; Hand and Uhlinger, 1992; Hand and Uhlinger, 1994; Stefanik et al., 2013).

In addition to a wealth of genomics data and functional genomics tools (reviewed in Röttinger, 2021), genome editing approaches have been used to generate several *N. vectensis* mutant and transgenic lines. To report tissue and cell type specific expression, initially promoters driving a fluorescent tag have been successfully integrated in the genome by random integration using I-SceI meganuclease (Renfer et al., 2010). Targeted genome editing has been achieved using TALEN as well as CRISPR/Cas9 (Ikmi et al., 2014). Since the successful use of CRISPR/Cas9 to generate *N. vectensis* knockouts and knock-ins (Ikmi et al., 2014), several other studies have used this versatile and easily adaptable tool to generate *N. vectensis* mutant lines. Nonetheless, the duration of the life cycle is crucial for enhancing the development of genetic tools in the *N. vectensis* community.

To grow *N. vectensis,* the most common feeding protocol is based on commercial stocks of *Artemia salina* (Decapoda) cysts that easily hatch into nauplii (Pan et al., 2022). During initial juvenile developmental phase of *N. Vectensis*, *Artemia* nauplii are too big to be fed to primary polyps, and smashed Artemia are used as source of food (Amiel et al., 2015). This protocol is also used in other cnidarians, e.g. *Clytia hemisphaerica* (Lechable et al., 2020). Following this initial phase, *N. vectensis* polyps are fed with freshly hatched *Artemia* nauplii until reaching sexual maturity. In addition, some laboratories are improving their feeding diet using pieces of oysters or mussels before inducing spawning (Hand and Uhlinger, 1992; Röttinger et al., 2012).

Although largely used as food for aquatic research models, it has been shown that *Artemia* cysts can be deleterious for the health of the fed organisms. In fact, they can be a source of pathogens, such as bacteria (*e.g., Mycobacterium sp., Vibrio sp*.) or chemical products (*e.g.,* arsenic) (Brix et al., 2003; Tkavc et al., 2011; Valverde et al., 2019). This source of contamination is further accentuated by using smashed *Artemia* nauplii as a source of food. This process creates a perfect incubation medium for bacterial growth and thus, increases the proliferation of unicellular organisms such as *Paramecia* that under dense conditions can be harmful to primary polyps (personal observations).

Rotifers (Gnathifera) have been successfully used in Zebrafish aquaculture allowing improved larvae growth and survival, in complement or in replacement of *Artemia* nauplii (Best et al., 2010; Lawrence, 2007). Rotifer cultures present many advantages such as a microscopic/small size (160-350μm), the possibility of dietary enrichment with nutrients/algae, a high proliferative rate, a small laboratory space occupancy, low levels of external pathogen contaminations (Lawrence et al., 2012) as well as less impact on the natural environment as we are maintaining the culture in the lab.

In this study, we describe optimized culture, rearing and gene-editing conditions, enabling the community working with the sea anemone to develop genetic tools in only a couple of months. We have adapted a semi-automatic culture system from zebrafish husbandry to maintain adult *N. vectensis*, and validated its capacity to facilitate the maintenance, the follow-up of the colonies and to improve the generation of offspring. Furthermore, we assessed the effects of rotifers, *Brachionus sp*. type « L », on *N. vectensis* juvenile polyp growth and identified the best culturing conditions (growth rate and survival), enabling to close the life cycle of males in 7 to 8 weeks and 12 to 13 weeks for females. As a reduced life cycle is a critical pre-requisite for developing genetic tools, we further indicate sgRNA design conditions that enable functional gene knockout by CRISPR/Cas9 using a single sgRNA as early as the F0 generation. Finally, we also show that CRISPR/Cas9 mediated knock-ins can efficiently be obtained both using homology direct repair (HDR) as well as non-homologous end-joining (NHEJ) compatible templates.

In sum, this work highlights new resources, *i.e.,* optimized husbandry and the rapid development of genetic tools, for the expanding scientific community studying *N. vectensis*. This improvement has the potential not only to increase scientific interest in this research model for EvoDevo questions but also for emerging topics such as environmental stress-response, whole body regeneration, and aging/longevity.

## Results

### Improved spawning efficiency using a semi-automated culture system

Although circulating culture systems are used in few laboratories working on *N. vectensis*, laboratory colonies are largely maintained in 250ml glass bowls containing ∼50 sexually mature polyps of *N. vectensis* in 1/3 ASW; pH 8,2; conductivity 15,5µS) (Figure 1D, Figure S1A). Adult polyps were maintained under constant dark conditions inside an incubator at 16°C (Figure 1D, Figure S2C) to avoid spontaneous spawning events, which could be induced by light and temperature stimulus (Fritzenwanker and Technau, 2002; Reuven et al., 2021). In these glass-bowls, 1/3 ASW is stagnant, and polyps were fed regularly (4 times per week with *Artemia* nauplii). To prevent any microorganism outburst and maintain *N. vectensis* polyps healthy, labor and time-intensive water changes were required. In consequence, this labor and time-demand increases proportionally with the size of the colony and number of genetic strains maintained in the laboratory.

To test whether we could reduce time spent for the daily animal routine in parallel to increase *N. vectensis* biomass, we adapted a semi-automated Zebrafish culture system for the specific needs of *N. vectensis* biology (Figure 1E, Figure S1D), *i.e.,* temperature, conductivity, water flux, water renewal. As adult polyps are maintained in dark condition using a light-blocking cabinet (Figure 1E, Figure S1D), the system for *N. vectensis* culture and maintenance was named “Zerolux”. We estimated that maintaining 13500 polyps (*i.e.,* max capacity of the Zerolux) using glass bowls in an incubator requires about 7 hours 30 minutes for cleaning and 5 hours 45 minutes for feeding, per week (Table 1). Differently, the same number of polyps in the Zerolux, required weekly about 3 hours for cleaning and 1 hours for feeding (Table 1). The implementation of a semi-automated culture system for *N. vectensis* represented therefore a reduction of 71% of the time invested in overall polyp maintenance.

**Table 1.**
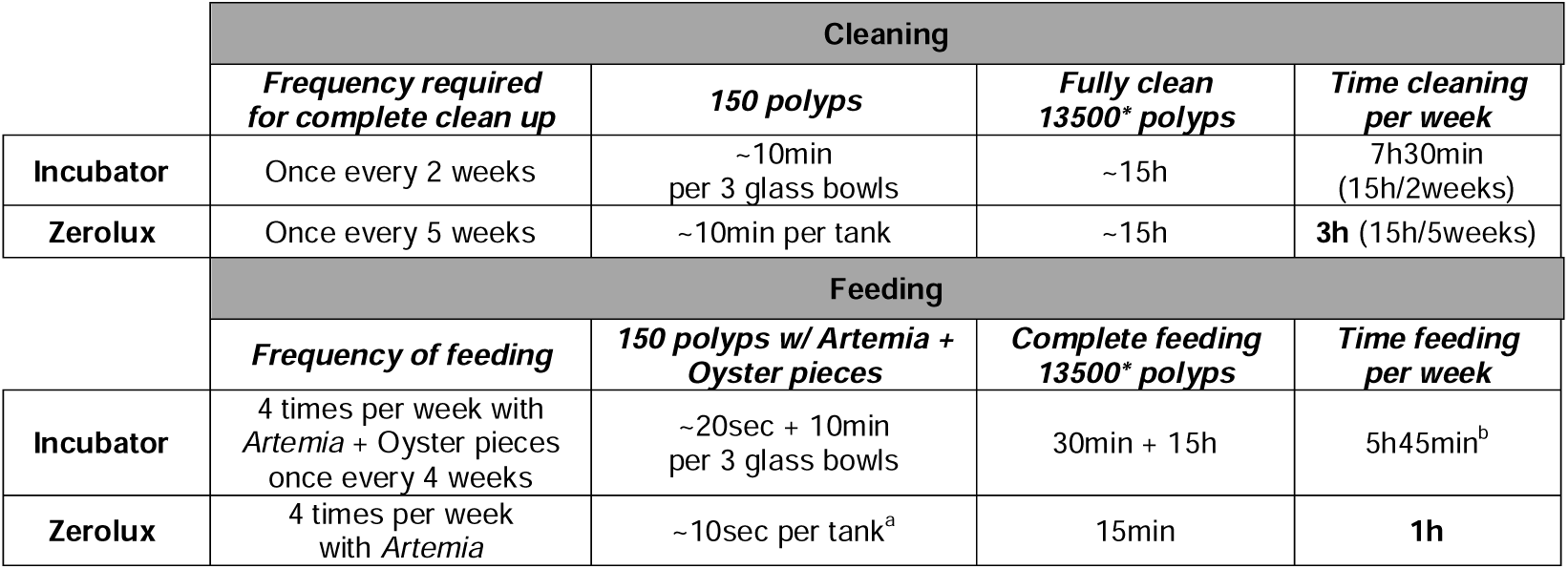
Estimation of time allocated to *Nematostella vectensis* maintenance tasks using glass bowls in an Incubator or the semi-automated system (Zerolux). *Zerolux max capacity (150 polyps x 90 tanks); ^a^ no Oyster pieces required; ^b^ 30min x 4 per week + 15h ÷ 4 (once every 4 weeks).

|  | Cleaning |  |  |  |
| --- | --- | --- | --- | --- |
|  | *Frequency required for complete clean up* | *150 polyps* | *Fully clean 13500\* polyps* | *Time cleaning per week* |
| **Incubator** | Once every 2 weeks | ~10min per 3 glass bowls | ~15h | 7h30min (15h/2weeks) |
| **Zerolux** | Once every 5 weeks | ~10min per tank | ~15h | 3h (15h/5weeks) |
|  | Feeding |  |  |  |
|  | *Frequency of feeding* | *150 polyps w/ Artemia + Oyster pieces* | *Complete feeding 13500\* polyps* | *Time feeding per week* |
| **Incubator** | 4 times per week with *Artemia* + Oyster pieces once every 4 weeks | ~20sec + 10min per 3 glass bowls | 30min + 15h | 5h45min<sup>b</sup> |
| **Zerolux** | 4 times per week with *Artemia* | ~10sec per tank<sup>a</sup> | 15min | 1h |

We then assessed whether the Zerolux system impacted *N. vectensis* spawning efficiency compared to glass bowls. To do so, we monitored sexually mature *N. vectensis* during 21 weeks for i) the female spawning induction rate (*i.e.,* number of females that spawned over the total of females stimulated to spawn, Figure 1F) and ii) estimated number of eggs spawned per female (Figure 1G). Four batches (*i.e.,* biological replicates) of 398 to 455 polyps were stimulated successively once every week – identified as *week 1, 2, 3* and *4*, respectively. These 4 independent replicates were monitored for a total of five spawning cycles: the 1^st^ cycle was done exclusively in glass bowls and occurred before part of the animals being transferred to the Zerolux. The next 4 full cycles of spawning were done in glass bowls as well as Zerolux tanks (*i.e.*, the polyps that were introduced in the Zerolux system).

The results show that the overall female spawning induction rate significantly increased in the Zerolux maintenance condition compared to the glass bowls, with median rates of 82% and 49.8% (Welch’s t-test p-value = 1.00e-08), respectively (Figure 1F). In addition, the number of eggs per female increased significantly in the egg clusters collected from the Zerolux maintenance condition in comparison to the glass bowls, with median values of 175 and 87 eggs/female (Welch’s t-test p-value = 0.0012), respectively (Figure 1G).

Overall, raising *N. vectensis* using a semi-automated culture system, such as the Zerolux, is less time-demanding, yields a more efficient spawning induction and increases the overall number of eggs produced by *N. vectensis* females.

### Feeding *N. vectensis* primary polyps with algal-enriched rotifers increases growth rate

After one week of embryonic development at 22°C, *N. vectensis* planula larvae develop/metamorphose into primary/juvenile polyps (Ormestad et al., 2011; Steger et al., 2022). With the current diet, *i.e.,* using smashed *Artemia* as first nutrition source once a week, it takes ∼4-6 months to reach sexually maturity in *N. vectensis* (Hand and Uhlinger, 1992). To optimize the growth rate of the juvenile polyps, we assessed the impact of multiple feeding diets on polyp growth (Figure 2A-C). From a batch of offspring that reached the primary polyp stage (spawned and fertilized in the same day), we selected 400 *N. vectensis* polyps with similar sizes for each condition. Primary polyps were cultured in glass bowls as they are too small for the Zerolux system.

**Figure 2.**
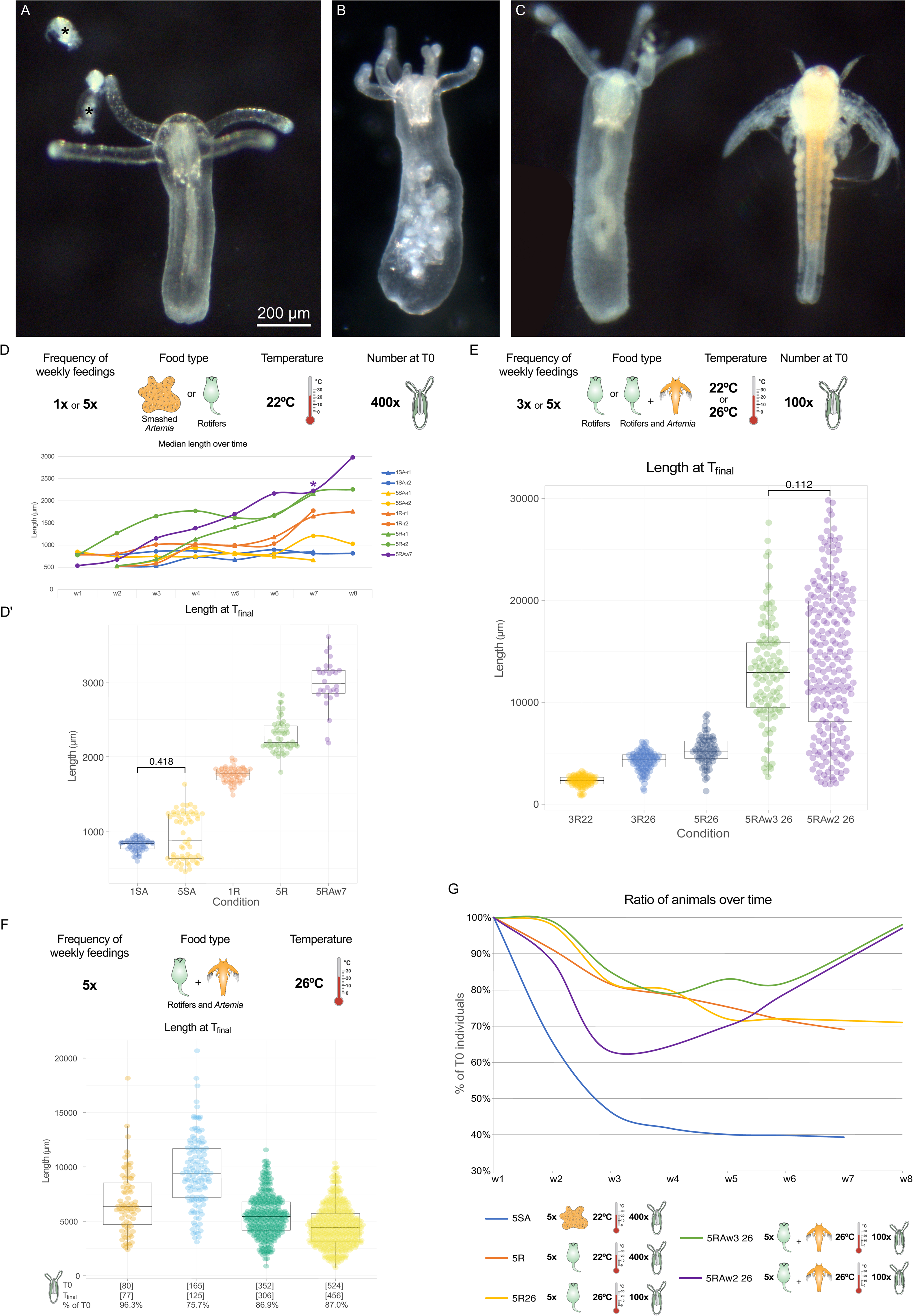
*Nematostella vectensis* body length growth can be optimized with a combination of live prey, feeding frequency, temperature, and polyp density. (A) 1 week-old *N. vectensis* juvenile polyp, in the presence of live Rotifers (indicated by the *). (B) 1 week-old *N. vectensis* juvenile polyp following Rotifers feeding. (C) 1 week-old *N. vectensis* juvenile polyp, in the presence of live *Artemia* nauplii. (D) Median body length of *N. vectensis* when comparing diets based on smashed *Artemia* nauplii or live Rotifers, given 1 or 5 times a week with an approximate initial density of 400 polyps, at 22°C, during up to 8 weeks. (D’) Detailed view of all the measurements made at Tfinal in the above conditions, by grouping each biological replicate. (E) Body length at Tfinal (8 weeks) of *N. vectensis* when comparing diets based on live Rotifers or live Rotifers followed by live *Artemia* nauplii, given 3 or 5 times a week with an approximate initial density of 100 polyps, at 22°C only (w0-w8) or 22°C (w0) followed by 26°C (w1-w8). (F) Body length of *N. vectensis* depending on the polyp density, for individuals fed 5 times per week, with live Rotifers and live *Artemia* nauplii (w3-w8), maintained at 22°C (w0) followed by 26°C (w1-w8) for 8 weeks. Numbers in the x axis represent, for each condition, the total number of polyps initially (T0) and at w8 (Tfinal), as well as the survival ration (% of T0). (G) Percentage of initial number of polyps present overtime for selected conditions described above. These values reflect the mortality, the polyps lost during maintenance of the culture (i.e. loss) as well polyps obtained by asexual reproduction (i.e. increasing). Post hoc Mann-Whitney U Test was used for pairwise comparisons amongst the independent conditions at Tfinal in A, B and C. Values are shown for non-significant results, otherwise p-value < 0.00001.

Over a period of eight weeks post fertilization, we monitored polyp size (30 randomly chosen polyps, measured between the tip of the mouth to the physa (Figure S2A-C)), depending on the applied feeding regime (smashed *Artemia salina* nauplii or live rotifers) and the frequency of feeding (1 to 5 times a week). The tested diets were: 1) **1** time per week **s**mashed ***A****rtemia* – 1SA; 2) **5** times per week **s**mashed ***A****rtemia* – 5SA; 3) **1** time per week live **r**otifers – 1R; 4) **5** times per week live **r**otifers – 5R; and 5) **5** times per week live **r**otifers then live ***A****rtemia* nauplii at **w**eek **7** (5RAW7) (Figure 2D, D’).

The initial median length of the polyps at T0 varied between 522μm and 808μm, depending on the condition (Figure 2D). In line with previous observations from the lab, the feeding regime with smashed *Artemia*, both 1 or 5 times per week led only to a mild increase in size during the 8 weeks of feeding, reaching a peak between 835μm and 870μm respectively at Tfinal (5 to 10% increase; Mann-Whitney U Test p-value 0.42) (Figure 2D, D’). Strikingly, our data showed that feeding with live algae-enriched rotifers largely and significantly increased the median body length of the polyps. In fact, at a feeding frequency of once a week (1R) the length increased 170% to 1769μm, and animals fed 5 times per week (5R) reached a median body length of 2192.5μm *i.e.,* 240% longer (Figure 2D, D’).

Interestingly we observed that polyps under the 1R or 5R diet reached a growth plateau after 7 weeks, suggesting that for larger polyps enriched rotifers may be too small to provide enough energy. Rather than increasing the rotifer concentration, we introduced live (un-smashed) *Artemia* nauplii at this specific phase (week 7, 5RAw7), and observed a new increase of the growth yielding a final median size 450% longer than T0, with a median body length at 2978.5μm (Figure 2D, D’). Thus, our results show that the use of live algae-enriched rotifers significantly increases the growth rate of *N. vectensis* during an initial phase (up to 7 weeks). Once the polyps become too large to be fed with rotifers, the introduction of live *Artemia* nauplii is sufficient to sustain polyp growth.

### *N. vectensis* growth is temperature and polyp density dependent

Growth in certain marine animals is temperature dependent (Angilletta et al., 2004). Thus, to test if *N. vectensis* juvenile polyp growth was also affected by the temperature, we tested the growth rate effects (batch size: 100 polyps, diet condition: **3** times rotifers per week) at two temperatures 22°C and 26°C (3R22 and 3R26, respectively) for 8 weeks (Tfinal). When comparing the median body sizes at Tfinal, we show that juvenile polyps almost doubled (84%) their median body length between 3R22 (2262μm) and 3R26(4166μm) (Figure 2E). These results indicate that temperature can significantly speed up the growing rate of *N. vectensis* juvenile polyps.

To go further, we tested if the addition of entire Artemia at an early stage (week 2 or week 3) may have an impact on the growth rate of the juveniles (Figure 2D, D’). At 26°C, we set up 3 different diets: 5R only, 5R complemented with live *Artemia* from week2 or week3 (5R26, 5RAw2 26 and 5RAw3 26, respectively). We observed that under these conditions the median length at Tfinal is further increased to 5166μm (128%), 12238μm (441%) and 14160μm (525%) for 5R, 5RAw3 and 5RAw2, respectively (Figure 2E).

Interestingly, in our initial set of experiments performed at 22°C we observed that feeding *N. vectensis* juvenile polyps using a 2R diet starting with 400 or 100 polyps per glass bowl (250ml), did not yield the same median body length at Tfinal, *i.e.,* 1989μm and 3581.5μm, respectively (Figure S2E). These data strongly suggest that polyp density may also affect polyp growth rate.

To test if polyp density plays indeed a role on the growth rate of *N. vectensis* juvenile, we set up a density experiment at 26°C using 4 different density conditions (1 - 80 polyps, 2 - 165 polyps, 3 - 352 polyps, and 4 - 524 polyps) using the best performing diet, *i.e.*, 5RAw3 (Figure 2F). At Tfinal the median *N. vectensis* polyp length was of 1) 6344μm, 2) 9417μm, 3) 5439μm and 4) 4426.5μm, respectively. Together with the results from the previous experiment that used a density of 100 (median polyp size Tfinal = 12238μm, Figure 2F) our results indicate that a density between 80-160 polyps/glass bowl, with an optimum of 100 polyps per glass bowl, at 26°C with the feeding diet 5RAw3 is best for a fast polyp growth. By assessing the number of polyps at the start (T0) and end (Tfinal) of the experiment, we were able to determine the loss ratio (number of polyps at Tfinal over initial number). Interestingly, the loss ratio for the four density conditions was of 0.037; 0.243; 0.131; and 0.13, respectively (Figure 2F), indicating that a density of 80-100 per glass bowl is ideal for polyp growth while preserving their survival.

In addition of following the *N. vectensis* juvenile polyps’ growth in various feeding diets (Figure 2D), temperatures (Figure 2E) and densities (Figure 2F), we also collected data regarding the polyp survival rates for the following conditions: 1) 5SA at 22°C, 2) 5R at 22°C, 3) 5R at 26°C, 4) 5RAw2 at 26°C and 5) 5RAw3 at 26°C (Figure 2G). Polyp survival rate was evaluated by counting the number of animals for each condition once per week. Our data show that 5SA at 22°C has the lowest survival rate with about 39% of the number of polyps at T0 still present at Tfinal. Strikingly, 5R at 22°C or 26°C led to a survival rate of 69% or 72% respectively, indicating the positive impact that rotifer have on the survival of the primary polyps. While the 5RAw3 26 condition performed like 5R22 and 5R26 during the first 4 weeks, we observed a poor survival rate for 5RAw2 26 during this same period (Figure 2G), indicating that re-introducing artemia to the feeding diet can have deleterious effects if performed too early, especially if the polyp size is too small to take them up. In fact, the 5RAw3 26 diet does not present such drastic decrease in polyp numbers during the first weeks (lowest point at 79%, Figure 2G). Unexpectedly, we observed that the “survival rates” (*i.e.,* number of polyps present in the glass bowl) for 5Rw2 26 and 5Rw3 26 increased again starting from week 3 and 4, for both diets respectively (Figure 2G), which could be associated with an increased physal pinching events (Figure S2F-H’’).

In sum, our results show that *N. vectensis* polyp growth and survival are positively affected by algae-enriched rotifers and higher temperatures and negatively affected by increasing polyp density, enabling us to determine 5RAw3 26 (**5** times algae-enriched rotifers per week, introduction of ***A****rtemia* nauplii at **w**eek **3**, polyp culture temperature **26**°C, culture density 100 polyps/glass bowl) as an optimal for fastest polyp growth.

### Optimized growing conditions speed up N. vectensis life cycle

*N. vectensis* is amenable to genetic manipulation. Thus, a reduced life cycle is crucial for the effective development of genetic tools. With the previous set of experiments, we validated the best diet for accelerating the growth of *N. vectensis* juvenile polyps: 5RAw3 at 26°C. Taking advantage of this optimized polyp growth condition, we assayed several stimuli to obtain sexually mature polyps to reduce the timing of the life cycle in *N. vectensis*. To do so, we set up a spawning assay for 13 weeks post-fertilization, with polyps raised using 5RAw3 26 (Figure 3). In sexually mature animal, *N. vectensis* spawning is induced following a 9h light and dT=4-6° temperature stimulus (Stefanik et al., 2013).

**Figure 3.**
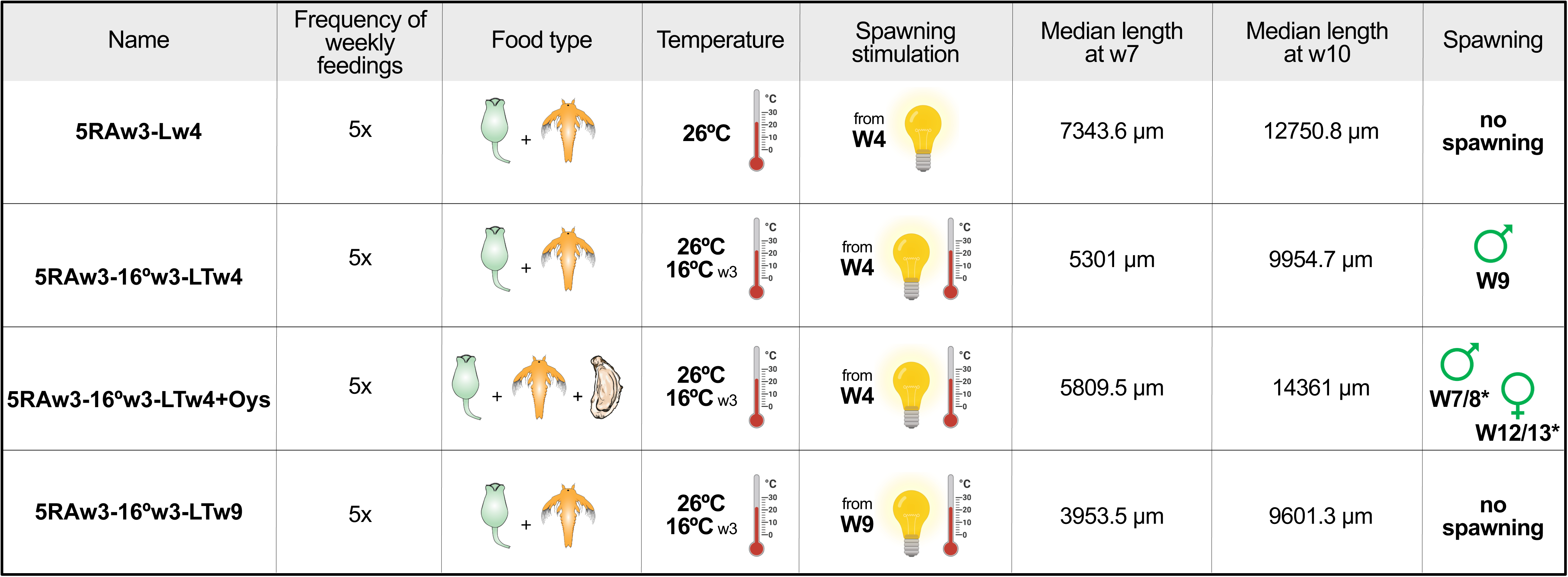
*Nematostella vectensis* spawning assays. Overview of the conditions used for to verify spawning success, during ??weeks. *N. vectensis* polyps were fed 5 times per week, with live prey (Rotifers or *Artemia* nuplii) and supplemented with Oyster for one of the conditions. The cultures were maintained either as 26°C or at 26°C (w1-w2) and then transferred at 16°C. The spawning stimulation occurred from the w4, every week. The median sized verified at week 7 and week 12 is shown for each condition, as well as the spawning success. Female and male symbols with the week number represent the first occurrence of female and male spawning under these 4 maintenance scenarios. * indicate that for different replicates the spawning week verified was not the same.

We therefore adjusted our culture condition accordingly by testing four set-ups: 1) **5RAw3** constantly at 26°C with weekly **l**ight stimulations starting at **w**eek **4** (5RAw3-Lw4); 2) **5RAw3** at 26°C for **3 w**eeks, then maintained at **16**°C with light and temperature (**LT**, 16->22°C) stimulations starting at **w**eek **4** (5RAw3-16°w3-LTw4); 3) **5RAw3** at 26°C for **3 w**eeks, then maintained at **16**°C with **LT** stimulation starting at week **4**, in which the animals were fed extra with **oys**ter fragments the day before spawning induction (5RAw3-16°w3-LTw4+Oys); and 4) **5RAw3** at 26°C for **3 w**eeks, then maintained at **16**°C with **LT** stimulation starting at **w**eek **9** (**5RAw3-16°w3-LTw9**) (Figure 3). We scored positive male spawning when the water containing stimulated *N. vectensis* was put in contact with non-fertilized eggs obtained from mature females, resulting in developing embryos. Female spawning was classified by the presence of eggs (Figure 3).

Interestingly, animals maintained at 26°C, light stimulated (*i.e.*, 5RAw3-Lw4) and monitored for 13 weeks never yielded any positive spawning events, neither females, nor males (Figure 3). Polyps under 5RAw3-16°w3-LTw9 conditions did not produce any gametes within the duration of the experiment neither. However, polyps released sperm and fertilized eggs following the 5RAw3-16°w3-LTw4 conditions as early as week 9 (Figure 3), indicating the importance of early LT stimulation for inducing sexual maturity. Nonetheless it is important to point out that following the same conditions (5RAw3-16°w3-LTw4), we did not observe any release of eggs within the timeframe of the experiment. Finally, the supplementation of the diet 5RAw3 with oyster fragments (5RAw3-16°w3-LTw4+Oys), further improved spawning efficiency. In fact, under these conditions the first male spawning was registered at week 7 and the first female spawning was observed at week 12 (Figure 3), suggesting that adding a rich source of nutrition, participates in the sexual maturation process of *N. vectensis*.

In sum, our results highlight that under a controlled, standard feeding regime and culture conditions, combined with LT stimulations during early polyp development, one can achieve sexual maturation and gamete production in *N. vectensis* in less than 2 or 3 months for males and females, respectively.

### CRISPR/Cas9 using one single-guide RNA can generate functional knockout animals at the F0 generation

Achieving to close the life cycle of *N. vectensis* in a relative short time frame (< 2 months), re-enforces the potential for the development of genetic tools. We therefore further explored the capacity to generate functional gene knockouts in *N. vectensis* using CRISPR/Cas9. Specifically, we aimed at optimizing the Cas9 sgRNAs pair to generate KO’s. For such, we took advantage of the already described natural red fluorescence present in the pharynx of polyps, which is encoded by a single gene, *fp-7r* (Ikmi and Gibson, 2010). We designed three new sgRNA’s specific targeting *fp-7r* and compared them to the originally described sgRNA used (Ikmi et al., 2014). We defined 3 regions of interest to design new sgRNA’s, based on an updated annotation for the gene, using RNAseq data (Johnston et al., 2021; Warner et al., 2018) (Figure 4A): 1) an intronic region upstream of the predicted transcription starting site for *fp-7r*, to be used as a non-coding cut site (*i.e.*, negative control), as potentially not affecting gene function, named sgR UTRi; 2) the first exon, corresponding to the same region previously targeted by CRISPR/Cas9, named sgR e1; and 3) the third exon, named sgR e3. We named the originally described sgRNA targeting exon 1 of *fp-7r* (Ikmi and Gibson, 2010), sgR Ex2*.

**Figure 4.**
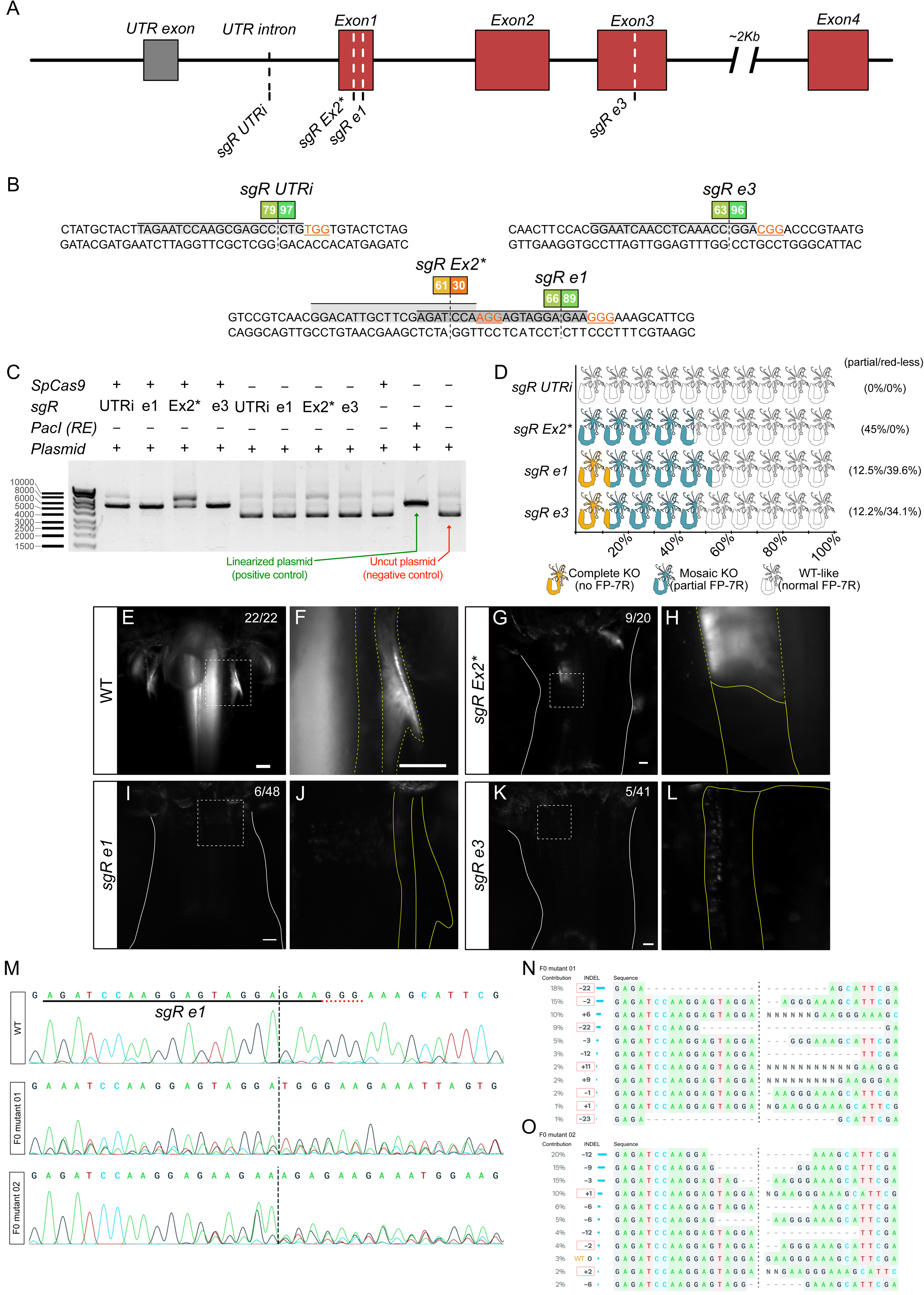
Generation of *Nematostella vectensis fp-7r* loss of function mutant. (A) Representation of the gene locus *fp-7r*. The 4 sgRNAs target regions are indicated by dashed lines. (B) Detailed overview of the targeted nucleotide sequence for each sgRNA (grey squares). The PAM is highlighted in orange and specific SpCas9 cut sites are indicated by dashed lines. Theoretical on-target score and off-target score determined by Benchling are indicated, with 100 being the best possible score. (C) SpCas9-sgRNA pair *in vitro* assay to validate DNA cutting capacity. (D) SpCas9-sgRNA complex *in vivo* assay results, showing the proportion of live adult polyps with 14 to 16 tentacles, showing partial or complete loss of FP-7R fluorescence. (E-L) Representative images of *N. vectensis* adult polyps FP-7R fluorescence for (E-F) wild-type controls; (G-H) partial ablated SpCas9-sgR Ex2 injected (I-J) red-less (i.e. knockout) SpCas9-sgR e1 injected; (K-L); red-less SpCas9-sgR e3 injected. Yellow doted lines highlight the position of actual FP-7R fluorescence, while solid lines represent the expected FP-7R fluorescence regions. (M) Sanger sequencing chromatograms for the wild-type control and two red-less animals obtained from SpCas9-sgR e1 injections. Best matching chromatogram sequence is shown, the position of the sgRNA e1 is shown with the clear indication of the expected cut site and the PAM in underlined by the red dotted line. (N-O) *In silico* decomposition of raw sanger results by ICE analysis (Synthego) and prediction of individual sequences obtained for (N) F0 red-less sgR e1 mutant 1 and (O) F0 red-less sgR e1 mutant 2. Abbreviations: UTR – untranslated region; sgR – sgRNA; KO – knockout. Scale bar: 100 μm. * indicates that this sgRNA was previously used in (Ikmi et al., 2014), therefore we used the original nomenclature, even though our current annotation of the gene suggest it locates in *fp-7r* exon 1.

For each region, we predicted several sgRNA using Benchling – Design CRISPR Guides tool, using the NCBI ASM20922v1 *Nematostella vectensis* genome version to obtain off-target scores. The best combination for higher in-target score (theoretical probability that a sgRNA induces Cas9 cut at that precise location), and higher off-target score (probability of no off-target existence in the genome) were selected (Hill et al., 2022). The in-target and off-target scores of the used sgRNAs are: sgR UTRi (79/97), sgR e1 (66/89), sgR Ex2 (61/30) and sgR e3 (63/96) (Figure 4B).

To determine if the obtained scores were potential indicators of DNA cleavage efficiencies, we first performed and *in vitro* assay for each of the above described sgRNAs in the presence or absence of *Sp*Cas9. As template, we used a plasmid containing a 2018bp region spanning the target regions. sgR e1 (66/89) and sgR e3 (63/96), which have comparable theoretical scores, were the most efficient. In fact, we observed a complete linearization of the plasmid compared to control conditions, *i.e.,* without *Sp*Cas9 (no linearization), or digested by PacI (linearization, Figure 4C). To our surprise, sgR UTRi that had the highest scores (79/97), only induced a partial linearization of the plasmid (Figure 4C). However, sgR Ex2 that displayed the lowest scores (61/30), performed poorly causing an even more partial linearization (Figure 4C). Thus, the theoretical scores alone are not indicative of the potential DNA cleavage efficiency. Further validation of sgRNA’s by an *in vitro* assay is crucial to define which sgRNA could be used for *in vivo* genome editing.

To evaluate to what extend the scores coupled with *in vitro* assays were indicative of a potential generation of functional KO’s, the 4 described sgRNA’s were co-injected with *Sp*Cas9 into zygotes. The knockout effects were evaluated by screening animals for the presence, partial-loss, or complete loss of FP-7R red fluorescence pattern in the 14-16 tentacle polyps (Figure 4D). As predicted by its target site, fertilized eggs, co-injected with *Sp*Cas9/sgR UTRi resulted into F0 polyps with a FP-7R fluorescence pattern, similar to WT animals (Figure 4D-F). Fertilized eggs co-injected with *Sp*Cas9/sgR Ex2, resulted into 45% (9 out of 20) F0 polyps with a partial loss of the FP-7R fluorescent pattern (9 out of 20 Figure 4D, G, H). Co-injection of *Sp*Cas9/sgR e1 or *Sp*Cas9/sgR e3 gave rise to both partial and complete fluorescence pattern loss. For *SpCas9*/sgR e1 12,5% (6 out of 48) and for *SpCas9*/sgR e3 12.2% (5 out of 41) F0 polyps completely lost their fluorescence pattern (Figure 4D, I-L), while partial loss of FP-7R fluorescence pattern occurred in 39.6% (19 out of 48) and 34.1% (14 out of 41), respectively (Figure 4D). These results suggest that the newly designed sgR e1 and sgR e3 induced, at a significant rate, a fully functional knockout of FP-7R already in the F0 generation. Both sgRNAs were also the ones that performed best in the *in-vitro* assay (Figure 4C).

Since *Sp*Cas9/sgR e1 co-injected animals yielded the most preeminent polyp number with a partial and complete loss of FP-7R pattern (Figure 4D), we further investigated the genetic nature of the mutations present in the F0 for two FP-7R functional KO polyps. By extracting genomic DNA from the physa region, followed by genotyping PCR (amplifying the region of interest) and Sanger sequencing, the results show a mix of several possible genotypes for the two mutants (Figure 4M). Further analyses of the sequences (deconvolution of the possible alleles using ICE by Synthego) revealed the potential INDELs caused by CRISPR. The combination of all predicted frameshift INDELs (Figure 4N, O) revealed that 46% and 16% of the INDELs gave rise to FP-7R KO’s, in mutant 01 and 02, respectively.

Our results reveal that for a given gene it is important to benchmark several sgRNA’s to achieve the highest/most complete gene knockout efficiency. They also show that using an *in silico* and *in vitro* efficient combination of sgRNA/Cas9 complex can yield to a fully functional knockout from the F0 generation, opening the possibilities of loss of function studies as early as in F0 polyps,

### Alternative genome insertion strategies can be used to generate knock-ins

While KOs in *N. vectensis* are becoming increasingly available, targeted knock-ins remain sparse (reviewed in Röttinger, 2021; Paix et al., 2023). To assay for CRISPR/Cas9-mediated targeted genomic insertion using the newly designed sgRNAs, we assessed if each individual *Sp*Cas9/sgRNA complex was able to generate knock-in mutants following a published protocol using HDR (Ikmi et al., 2014). In this series of experiments, via the combination of sgRNA location and repairing template we also assayed whether genomic insertion NHEJ can be achieved in *N. vectensis* (Figure S3). In fact, the sgRNAs are located 353bp and 854bp (sgR UTRi and sgR e3, respectively) away from the uncut homology arm (Figure S3C, D), greatly reducing the possibility (< 80%) of genomic insertions by HDR events (Elliott et al., 1998; Zhang et al., 2017), and thus allowing the possibility to score NHEJ-mediated knock-ins.

To do so, we co-injected a single *Sp*Cas9/sgRNA complex with a donor vector that contains one kilobase of left and right genomic homology arms flanking the *fp-7r* exon 1 (Figure 5A). The donor vector contains a reporter gene, *mCherry,* under the control of the *N. vectensis myosin heavy chain1* (*MHC1)* promoter (Figure 5B). Thus, successful integration resulted in mCherry expression in the retractor muscles (Renfer et al., 2010). When using *Sp*Cas9/sgR Ex2 or *Sp*Cas9/sgR e1 injection (Figure S3B), we observed a clear mosaic mCherry expression associated with muscle cells upon successful homologous recombination occurring in *fp-7r* exon 1 of the F0 lines. *Sp*Cas9/sgR UTRi or *Sp*Cas9/sgR e3 complexes induced Cas9 mediated cleavage both in the genomic locus of interest as well as in the donor vector (Figure S3C, D). As the distance between sgRNA site and the region of the repairing template strongly reduce any HDR-mediated insertions (Elliott et al., 1998; Zhang et al., 2017), the observed genomic insertion of mCherry in *fp-7r* UTR or exon 3 (muscular staining, Figure 5) are most likely mediated by NHEJ.

**Figure 5.**
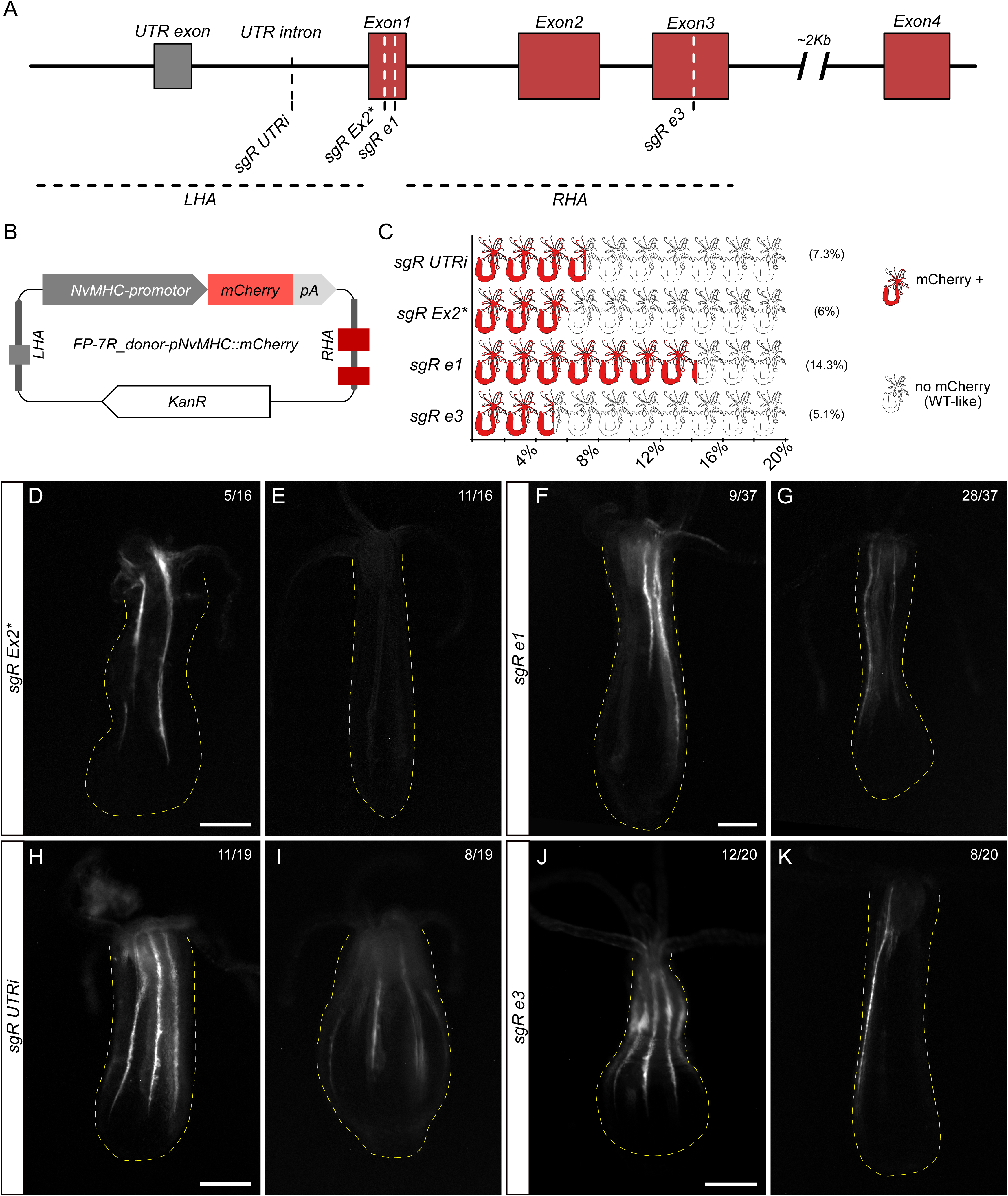
Generation of *Nematostella vectensis* knock-in at the *fp-7r* gene locus. (A) Representation of the gene locus *NvFP-7R*. The 4 sgRNAs target regions are indicated by dashed lines as well as the regions used as repairing template, the left homology arm, LHA, and the right homology arm, RHA. (B) Overview of the repairing plasmid used to *in vivo* validate knock-ins. (C) SpCas9-sgRNA complex *in vivo* assay for knock-in, showing the proportion of live polyps, showing NvMHC1 driven mCherry fluorescence. (D-K) Representative images of *N. vectensis* polyps showing NvMHC1 driven mCherry fluorescence in (D-E) SpCas9-sgR Ex2 injected; (F-G) SpCas9-sgR e1 injected; (H-I) SpCas9-sgR UTRi injected; and (J-K) SpCas9-sgR e3 injected. Yellow dotted lines show the delimitation of the body of *N. vectensis* polyps. Scale bar: 100 μm. * indicates that this sgRNA was previously used in (Ikmi et al., 2014), therefore we used the original nomenclature, even though our current annotation of the gene suggest it locates in *fp-7r* exon 1.

To evaluate *Sp*Cas9/sgRNA mediated insertion efficiency we scored animals for muscular mCherry fluorescence at 4 weeks post fertilization (6-8 tentacles), timing for which none of the negative controls (*i.e.*, embryos injected only with the donor vector without *Sp*Cas9/sgRNA) showed fluorescence. When comparing *Sp*Cas9/sgRNAs pairs that could mediate HDR insertion, we observed that following *Sp*Cas9/sgR Ex2 injections, 6% (16 out of 267) displayed mCherry expression at the expected location in the retractor muscles (Figure 5C, D, E). *Sp*Cas9/sgR e1 injections were twice more efficient, with14.3% (37 out of 259) of polyps showing mCherry localized expression in the retractor muscles (Figure 5C, F, G). Compellingly, when using the two *Sp*Cas9/sgRNAs pairs that are not localized in the exon 1 (sgR UTRi & sgR e3) we also observed mosaic expression of mCherry in the retractor muscles. In fact, 7.3% (19 out of 259) or 5.1% (20 out of 389) of the analyzed polyps expressed mCherry in the expected tissues after *Sp*Cas9/sgR UTRi or *Sp*Cas9/sgR e3 injection, respectively (Figure 5C, H-K). Based on the homology arms configuration and following *Sp*Cas9 DNA cleavage both in the genomic DNA locus as well as in the repairing template (Figure S3) these results suggest that non-homologous end joining DNA repair occurred. Of note, amongst the animals that expressed mCherry in the retractor muscles, one could observe several degrees of mosaicism ranging from a few patches of fluorescent cells (low occurrence, Figure 5E, K) to polyps that express mCherry in almost all retractor muscles (high occurrence, Figure 5H, J).

In sum, our data in *N. vectensis* show that the verified (*in silico*, *in vitro* and *in vivo*) efficiency of sgRNA to generate KO’s correlates with insertion occurrence. Furthermore, not only HDR-, but also NHEJ-mediated insertion can be used to generate stable knock-in lines, in which NHEJ process seems to generate less mosaic profiles.

## Discussion

In this study we develop complete and detailed community resource for the generation of offspring biomass, fast growing and improvement of CRISPR/Cas9 mediated knockout and knock-in for *N. vectensis*. With the comparison of multiple maintenance and feeding conditions, we highlight maintenance conditions leading to an increased sexual reproduction rate, body length growth and offspring survival. We reduce the life cycle of *N. vectensis* from 4-6 months to 2 months. This is a great advantage for the development of genetic strains in this research model. By comparing the efficiencies of a set of *Sp*Cas9/sgRNAs complexes, which target a single gene (*fp-7r*) for the establishment of KO and KI lines, we reveal that a combination of *in silico* prediction and *in vitro* validation of the sgRNAs is an efficiency indicator for frame shifting INDELs. We further show that in addition of HDR mediated insertion, NHEJ-mediated insertion can be envisioned for the establishment of targeted KI lines.

Maintaining only a few dozen adult *N. vectensis* in the laboratory, which involves keeping them in glass bowls with stagnant water, changing the water weekly or bi-weekly, and feeding them three to four times a week with *Artemia* nauplii, (Al-Shaer et al., 2021; Hand and Uhlinger, 1992) may not be time-consuming, although this may have an effective cost in terms of spawning induction in sexually adult polyps (Genikhovich and Technau, 2009). However, when scaling the *N. vectensis* colony for high polyp density, the maintenance will then become time consuming task. Here we show that an alternative method to maintain *N. vectensis* using a semi-automatic modified fish aquaculture system can be effectively used. This semi-automated aquarium system reduces the amount of time needed to maintain a *N. vectensis* adult culture, with the benefit of increasing both the fertility rate and the overall gamete production (*i.e.*, eggs). Thus, this system could be an alternative to maintain, under optimized conditions, a very large population of *N. vectensis*, as well as it could be a great tool to maximize the reproduction of a rather small and restricted population, as for example a specific *N. vectensis* transgenic line.

In the wild, *N. vectensis* adult polyps are a few centimeters in length and will reach sexual maturity in approximately six months or less (Williams, 1983). In this study, by measuring systematically the length of the body (from the base of the tentacle crown to the physa) of *N. vectensis*, we revealed that growing could be accelerated using specific nutrient conditions and temperature increase (Amiel et al., 2015; Ormestad et al., 2011). Although, due to nature of *N. vectensis* predation technique (*i.e.*, passive predator that captures passing prey), we revealed that depending on the stage of the life cycle (*i.e.,* size), different kinds of prey are required for an optimal growing. Here we tested several combinations of live prey, in contrast with the previous established method exclusively based on *Artemia salina* nauplii (smashed for early polyps and live for bigger animals) (Amiel et al., 2015). By introducing at the early polyp stage live algae-enriched Rotifers as a prey, we largely improved the growth speed and simultaneously reduced the mortality rate. This kind of approach combining live prey from diverse sizes (*e.g.*, Rotifers and *Artemia),* adapted to the size of the animal life stage, is widely used in aquaculture (Dhont et al., 2013; Pan et al., 2022) as well as for established animal models such as zebrafish (Dabrowski and Miller, 2018).

During our observation for over two months of several feeding conditions, it is important to mention that the feeding regimes with early introduction of live *Artemia* nauplii (*i.e.,* at week 2 or week3) generated a highly variable body length distribution, 1966μm to 29817μm and 2673μm to 27626μm respectively (Figure 2E). This variability can be attributed to a combination of two factors: (1) at these early stages (w2 or w3), not all *N. vectensis* juvenile polyps will be able to promptly feed on live *Artemia*. Thus, slightly bigger polyps will grow faster in early time points, and therefore smaller polyps will have a delayed growth compared to the bigger ones; (2) a limitation of the experimental design, in which all polyps present in the dishes were considered without systematically excluding newly pinched polyps (asexual reproduction) (Al-Shaer et al., 2023; Reitzel et al., 2007). Supporting this hypothesis, depending on the conditions, we observed that a high number of *N. vectensis* polyps undergoes asexual reproduction by physal pinching. This may be due to the very rich and intense diet in which these polyps are maintained, suggesting that physal pinching asexual mode of reproduction can be trigger by highly nutritive conditions.

The recent advent of CRISPR/Cas9 (Doudna and Charpentier, 2014) has revealed a promising genome editing tool for relatively simple generation of gene specific knockout and insertion of exogenous DNA, *i.e.,* knock-in (Momose and Concordet, 2016). The first CRISPR/Cas9 knockouts and knock-ins in *N. vectensis* have been demonstrated since a few years (Ikmi et al., 2014). Several studies have been addressing from a functional point of view the generation of knockouts in *N. vectensis*, most of these studies rely on the fertilized oocyte injection of two or several sgRNA’s in parallel to drive CRISPR/Cas9 mutagenesis (Babonis et al., 2021; Chen et al., 2020; He et al., 2018; Ikmi et al., 2020; Nakanishi and Martindale, 2018; Servetnick et al., 2017; Wijesena et al., 2017; Zang and Nakanishi, 2020). Our study reveals the possibility of generating CRISPR/Cas9 *N. vectensis* mutants using a single guide RNA (Ehrlich et al., 2022; Gahan et al., 2022; Kraus et al., 2016; Tournière et al., 2020) each designed at different location in the coding region, thus reducing the risk of off-site targets (Höijer et al., 2022).

Although by targeting an endogenously expressed fluorescent protein (FP-7R) we also revealed that functional deletion of a given gene can be obtained in the entire animal as early as the F0 generation in *N. vectensis*. Even if these results can be gene specific, it reveals a very promising road in which one can study, directly from the F0 generation, the involvement of specific genes underlying complex biological key processes such as sexual or asexual reproduction (*e.g.*, gonad maturation, spawning), environmental stress-response, regeneration, longevity or other essential adult-specific processes (*e.g.*, homeostasis, aging). From a genetic point of view, the need to further generate filial generations (e.g., F1, F2) is important to purify and stabilize a single inactivating mutation. Although some genes of interest might induce severe phenotypes, leading to, for instance, embryonic lethality and infertility. Thus, a F0 KO could be an effective way to circumvent these extreme phenotypes since *N. vectensis* can be clonally (asexually) expanded.

In *N. vectensis* the genomic insertion of genetic material has been classically and largely based on the I-SceI (intron-encoded endonuclease I from *Saccharomyces cerevisiae*) – meganuclease approach (Admoni et al., 2020; Busengdal and Rentzsch, 2017; Havrilak et al., 2017; He et al., 2018; Ikmi et al., 2020; Nakanishi et al., 2012; Renfer et al., 2010; Richards and Rentzsch, 2014; Steinmetz et al., 2017; Sunagar et al., 2018; Tournière et al., 2020). I-SceI protein stimulates homology recombination (HR) by creating a site-specific double strand breaks (DSB) in the genome, in a process called homing (Chames et al., 2005; Silva et al., 2011). Although, CRISPR/Cas9 has also been used to generate genomic insertions in *N. vectensis* with long homology arms (Ikmi et al., 2014), more recently, a study showed increased efficiency using short homology arms (Paix et al., 2023). These insertions are mainly of two types: large regulatory regions located upstream of coding DNA (i.e. “promoter”) followed by a reporter fluorescent protein (He et al., 2018; Ikmi et al., 2014) and gene specific insertion of a reporter fluorescent protein (Gahan et al., 2022; Lebedeva et al., 2022; Lebouvier et al., 2022; Paix et al., 2023). Interestingly, all these CRISPR/Cas9 mediated knock-ins have been generated using site-specific HR but using two alternative strategies based on long homology arms, about 1Kb long (He et al., 2018; Ikmi et al., 2014; Lebedeva et al., 2022; Lebouvier et al., 2022), or short homology arms, 40-50bp (Gahan et al., 2022; Paix et al., 2023; Seleit et al., 2021). Nevertheless, our current study show that similarly to axolotl (Fei et al., 2018), there is also the possibility to generate CRISPR/Cas9 genomic insertions using NHEJ (Figure S3, Figure 5). NHEJ is error-prone event in which DNA broken ends are joined together (Miyaoka et al., 2016) while HDR is a precise DSB repair mechanism using homologous recombination. Although, HDR requires that the homology repairing template locates near the cut site, since 100-200bp away from the cut reduces KI efficiency of 80% (Elliott et al., 1998; Zhang et al., 2017).

In our study, we designed an approach using a single plasmid to test both HDR mediated genomic insertion as well as NHEJ. Our results revealed that CRISPR/Cas9 HDR mediated insertion generated a higher number of positive transgenes when compared to NHEJ, although showing a wide variability of mutated regions (*i.e.*, mosaic expression). In contrast, the lower number of knock-ins obtained by NHEJ showed a large spectrum of transgene expression. The difference between the natural occurrence of these two mechanisms of DSB repair and how these work in *N. vectensis* is largely unknown. Interestingly, in mammalian cells DNA DSBs are predominantly repaired by NHEJ (Chiruvella et al., 2013). In contrast, other organisms such as yeast favor homologous sequences to repair DNA breaks (Lieber, 2010). Furthermore, the occurrence of DNA repair by either mechanism has been shown to be cell type and gene locus specific under Cas9 DSB experiments (Miyaoka et al., 2016). More recently, in mice and rat, encouraging results have been obtained by driving genome insertion under a setup allowing both repairing mechanisms to operate (Yoshimi et al., 2021). The generation of *N. vectensis* CRISPR/Cas9 knock-in is at its early age and our results reveal that considering NHEJ as an approach for genomic insertion could represent an alternative to HDR or even a tool to apply in parallel.

Our study further brings to light the genetic engineering interest of *N. vectensis*, and thus its value to address various biological questions, ranging from evolution, embryonic development, regeneration to stress-response and aging. The optimized and robust life cycle improvement here described puts *N. vectensis* generation time at the level of widely used genetic models such as zebrafish and mice (∼2 months) (Lieschke and Currie, 2007). We show that full functional knockouts and wide genomic integration can be obtained as early as the F0 generation. Combined with the extreme capacity to regenerate and long-living features of *N. vectensis* (Amiel et al., 2021; Röttinger, 2021), this can be a great tool to rapidly expand a population of mutants. In sum, we reveal an optimized and time efficient method to maintain, grow, reproduce, and generate mutants (both knockouts and knock-ins) of *N. vectensis*.

## Material and Methods

### Maintenance and spawning induction of N. vectensis colonies

*N. vectensis* adults, juveniles and embryos were cultivated according to (Amiel et al., 2015; Amiel et al., 2019; Röttinger et al., 2012). Animals were kept in glass bowls (250 ml) with 1/3x artificial seawater (1/3ASW) (salinity: 12pp; PRODIBIO Pure Ocean - Expert Reef Salt) (Figure S1). To keep the animals in a healthy reproductive state, they were kept in an incubator at 16°C in the dark and water was changed once a week (Figure 1D, FigureS1C). Adult animals were fed three times a week with *Artemia* nauplii and with oyster’s fragments one day prior to spawning. Female and male spawning were induced by a 9h light and temperature stimulus, followed by water changes to synchronize the spawning events. Oocytes were collected. The gelatinous mass around the eggs was removed with 2–4% L-Cysteine in 1/3ASW before fertilization and then washed 3 times within 1/3ASW. For a simultaneous development of the embryos, all oocytes were fertilized in glass dishes at the same time, by adding water containing *N. vectensis* sperm. The embryos were kept in dark, in filtered 1/3ASW at 22°C, until the desired stage. In the present series of experiments, metamorphosed polyps were manually sorted out by overall similar morphology (size, tentacle number and length) and placed in glass bowls in batches of determined number of individuals for each of the experimental condition.

### Artemia salina and *Rotifers culture*

5ml of commercial *Artemia salina* cyst (Ocean Nutrition, Planktovie, Marseille France) were added in 1L of 1/3ASW. Live artemia nauplii were collected after 48h of incubation at 22°C. Artemia were then cleaned from the remaining cysts/debris, placed in clean 1/3ASW and use to fed *N. vectensis* polyp colonies.

Rotifers were cultivated according to (Lawrence et al., 2012) and following Planktovie (Marseille, France) protocol : https://planktovie.biz/protocoles/protocole-de-culture-du-rotifere/). Compact Culture System (CCS) of 14L 1/3ASW was installed. The inoculation of the culture was initiated by adding 1million L-type rotifers in the CSS bucket. Rotifer culture was continuously oxygenated with an air pump and maintained at constant 25°C with the help of an immersible heater. To avoid the accumulation of detritus in the Rotifers culture, the entire CCS system was clean once a week, while the floss was rinsed every two days. The CCS was link to a peristaltic pump to provide mixed algae enrichment to the rotifer culture (RG complete, Reed MariCulture Inc., Campbell, CA, USA). Continuous feeding of the rotifers with RG complete diluted 50 times in 1/3ASW was placed in a dedicated container (2,5L) and was dispensed over the course of the day to the Rotifer culture bucket by the peristaltic pump (600ml per 24h). The microalgae solution (RG complete) was kept in suspension using a bubbler to avoid sedimentation and clogging of the tubing.

Rotifers concentration was monitored once a week by mixing 0,5 ml of the Rotifers culture within 0,5ml of white vinegar (i.e. to stop rotifers swimming behavior). The number of Rotifers was determined by counting the total number of individuals within the 1 ml mixture, using a cell counter chamber. Counting was repeated three times and the mean value used to determine the rotifers concentration in the culture. Gravid Rotifers (visible white eggs were observable in the posterior region of the animlas) were also counted regularly as indicator of the dynamic growth of Rotifers culture.

To keep a fast growth of the colony, 4L of the Rotifer culture were harvested every two days. The 4L were filtered using filters from two specific mesh size: 0,12mm to remove algae clogs and 0,041mm to collect the rotifers. Rotifers were re-suspended in 1/3ASW to a final concentration of 10000 rotifers/ml as a working feeding solution. 1/3ASW was added into the CCS to maintain a total volume of 14L in the Rotifers culture.

### Zerolux technology and monitoring of the physico-chemical parameters

The Zerolux system is an aquatic husbandry technology adapted from the zebrafish husbandry rack ZebTec from Techniplast S.p.A., Buguggiate VA, Italy. This technology allowed to maintain several aquariums organized along lines (6 letters from A to F) and rows (15 numbers from 1 to 15). A total number of 90 aquariums, 1,1L each, were maintained into the Zerolux conditions (Figure S1D). A dark cabinet was adapted around the ZebTec to maintanin the *N. vectensis* colonies into dark conditions, thus the name Zerolux. In addition, we adapted the default water solutions flow to the 1/3ASW (pH8,2; conductivity 15,5µS) necessary to raise *N. vectensis* (Figure 1E, Figure S1D). Automatic balance of the salinity was made by incorporation of reverse osmosis water (RO water) when needed. A total of 3% to 5% of 1/3ASW from the entire system were changed automatically every 24h in a progressive manner. The 1/3ASW flow is constantly running and filtered through mechanical (100um mesh size and UV) and chemical (charcoal) filters.

### Single guide RNA design and synthesis

Benchling tool “CRISPR Design and analyze guides” (https://benchling.com) was used to design single guide RNAs (sgRNAs) to target different regions of the *fp-7r* gene (Ikmi and Gibson, 2010). Upon the analysis of the CDS region for *fp-7r* previously described, and compared to recent transcriptomic (Warner et al., 2018) and genomic resources (Zimmermann et al., 2022) 3 regions of interest were identified: an intronic region upstream of the predicted *fp-7r* start codon; the first exon, corresponding to the same region previously targeted by CRISPR/Cas9; and the third exon of the *fp-7r* gene. We identified 1 sgRNA sequence for each region: sgR UTRi (5’ TAGAATCCAAGCGAGCCCTG<u>TGG</u> 3’), sgR e1 (5’ AGATCCAAGGAGTAGGAGAA<u>GGG</u> 3’), and sgR e3 (5’ GGAATCAACCTCAAACCGGA<u>CGG</u> 3’).

sgRNA synthesis was caried out used a cloning-free method (Varshney et al., 2016) based on oligos. 4 target-specific DNA oligos which contain a 17-nt-long T7 promoter + GG, followed by 18-nt target sequence (18 nucleotides 5’ of the protospacer adjacent motif – PAM) and a 20-nt sequence complementary to the guide RNA were ordered. A “generic” DNA 80-nt long oligo for the guide RNA region was further ordered. The two oligos were annealed (at equimolar concentrations 10 μM each) and extended with Phusion DNA polymerase (Thermo Fisher Scientific, Illkirch-Graffenstaden, France), and the resulting product was used as a template for in vitro transcription. sgRNA synthesis was carried out using NEB T7 High Yield RNA Synthesis kit (New England Biolabs France Genopole, Evry, France), following manufacturer’s instructions. The final product was purified using Zymo Research RNA Clean & Concentrator -5 kit (Zymo Research Europe GmbH, Freiburg, Germany) and it was run in 1.5% agarose gel to verify successful sgRNA synthesis and quality.

### Transgenic microinjection settings

Injections into fertilized eggs for knockout experiments were done using 50ng/µl of each sgRNA independently and 80nM of NEB EnGen Spy Cas9-NLS protein (New England Biolabs France Genopole, Evry, France), pre-incubated at room-temperature for 15 minutes (Hill et al., 2022). The injection mix was supplemented with 20ng/µl of Cas9-NLS mRNA (Addgene #141108) and 50Cng/μl of Invitrogen Dextran, Alexa Fluor 568 (Thermo Fisher Scientific, Illkirch-Graffenstaden, France). For knock-in, the mix was prepared as described above and 30ng/µl of the plasmid with the repairing template was further included in the mix.

### Imaging

Live polyps (wild type and transgenic) were imaged after a relaxation period of 15min in 7,14% in MgCl_2_ + 1/3ASW on the light table. The imaging setup was composed of either with a Zeiss Stereo Discovery V8 or a Zeiss Axio Imager A2 (both Carl Zeiss Microscopy GmbH, Jena, Germany) equipped with a Canon 6D digital camera, triggering two external Canon Speedlite 430 EX II Flashes and controlled by the Canon Digital Photo Professional software (Canon Inc., Tokyo, Japan) or a Zeiss Axiocam 506 color camera running with the ZEN 2009 software (Carl Zeiss Microscopy GmbH, Jena, Germany), respectively. ImageJ (Schneider et al., 2012) segmented line followed by measurement tool was used to determine the length of *N. vectensis* polyps (Figure S2). Images were edited using Photoshop CS6 software (Adobe Systems Inc., San Jose, CA, USA) as well as Affinity Photo (Serif Europe Ltd, West Bridgford, United Kingdom) and compiled together using Affinity Design (Serif Europe Ltd, West Bridgford, United Kingdom).

## Acknowledgments

The authors thank Kevin Foucher for initial observations on the optimization of the feeding regime, the IRCAN Marine Invertebrate Facility for (anti)Aging Research (ANTIAGE), as well as Valérie Carlin, Brigitte Poderini, Thamilla Zamoum, and Renaud Rebillard for animal care. We thank past and current team members for fruitful discussions and Matthew Gibson for the plasmid containing *N. vectensis* codon optimized *Sp*Cas9-2xNLS (addgene # 141108).

## Data availability

All relevant data can be found within the article and its supplementary information.

---

**Figure S1 – *Nematostella vectensis* maintenance conditions.** (A) *N. vectensis* sexually mature polyps in 250 mL glass bowls. (B) *N. vectensis* sexually mature polyps in 1.1L Zerolux tanks. (C) Organization of *N. vectensis* colonies inside the incubator. (D) Organization of *N. vectensis* colonies in the Zerolux system.

**Figure S2 – *Nematostella vectensis* length measurement set up, results and asexual reproduction.** (A) Adult polyp of *N. vectensis* with yellow dotted line indicating the procedure to obtain the length. (B) Representative picture obtained prior to measurement of *N. vectensis* polyps. (C) Detailed view showing the lines (in black) used to measure *N. vectensis* polyps. (D) Median body length of *N. vectensis* when comparing diets based on live Rotifers or live Rotifers followed by live *Artemia* nauplii, given 3 or 5 times a week with an approximate initial density of 100 polyps, at 22°C only (w0-w8) or 22°C (w0) followed by 26°C (w1-w8), during up to 8 weeks. Violet and green asterisks (*) indicates when live *Artemia* were introduced in the diet. (E) Initial observation of *N. vectensis* length at Tfinal (week 8) for differential growth depending on the density (100 polyps vs 400polyps at T0), under a diet of live Rotifers twice a week at 22°C. (F-H’’) Pictures of asexual reproduction of *N. vectensis* obtained during the follow up of multiple growing diets. (F) An example of the initial deformation in the body plan of *N. vectensis*, showing an early stage of asexual reproduction. (G) An example of an almost complete physal pinching occurring. (H-H’’) Three examples of newly pinched polyps with different levels of tentacles development. (I) Body length of *N. vectensis* at w7 and w10 for each of the 4 feeding and maintenance conditions used for the spawning assays detailed in Figure 3.

**Figure S3 – Schematic representation of HDR- or NHEJ-mediated insertion.** (A) Schematic representation of non-homologous end joining (NHEJ) and homology direct repair (HDR), following double strand breaks induced by Cas9. (B) Overview of HDR mediated insertion tested. Scheme for the initial DNA templates (upper panel), i.e. genomic DNA at the *fp-7r* locus and plasmid containing the repairing template flanking NvMHC::mCherry. In the case of using *Sp*Cas9 coupled with sgR Ex2 or sgR e1, only the genomic DNA will be cleaved by *Sp*Cas9 (middle panel). Thus, double strand breaks repaired via HDR, using the plasmid as template, will originate a scenario with NvMHC::mCherry inserted at the *fp-7r* exon 1 region (lower panel). (C) Overview of NHEJ mediated insertion tested in the *fp-7r* UTR region. Scheme for the initial DNA templates (upper panel), i.e. genomic DNA at the *fp-7r* locus and plasmid containing the repairing template flanking NvMHC::mCherry. In the case of using *Sp*Cas9 coupled with sgR UTRi, both the genomic DNA and the repairing plasmid will be cleaved by *Sp*Cas9 (middle panel). Thus, double strand breaks repaired via NHEJ will connect both cut ends, originating a scenario with NvMHC::mCherry inserted in the *fp-7r* UTRi region (lower panel). (D) Overview of NHEJ mediated insertion tested in the *fp-7r* exon 3. Scheme for the initial DNA templates (upper panel), i.e. genomic DNA at the *fp-7r* locus and plasmid containing the repairing template flanking NvMHC::mCherry. In the case of using *Sp*Cas9 coupled with sgR e3, both the genomic DNA and the repairing plasmid will be cleaved by *Sp*Cas9 (middle panel). Thus, double strand breaks repaired via NHEJ will connect both cut ends, originating a scenario with NvMHC::mCherry inserted in the *fp-7r* UTRi region (lower panel).Dashed yellow and orange lines represent the left and the right homology arm, respectively. The location of the sgR position is shown by an arrowhead. The representations for genomic DNA and plasmid DNA are not scaled.

**File S1 – 1 week-old *Nematostella vectensis* polyp first feeding with live Rotifers.** The video shows the first feeding event (with live Rotifers) of 1 week-old *N. vectensis* juvenile. The video was recorded for a total of 13 min, thus here it runs 11X faster than real time.

## References

Admoni, Y., Kozlovski, I., Lewandowska, M. and Moran, Y. (2020). TATA Binding Protein (TBP) promoter drives ubiquitous expression of marker transgene in the adult sea anemone *Nematostella vectensis*. Genes 11, 1081.

Al-Shaer, L., Havrilak, J. and Layden, M. J. (2021). *Nematostella vectensis* as a model system. In Handbook of Marine Model Organisms in Experimental Biology, p. CRC Press.

Al-Shaer, L., Leach, W., Baban, N., Yagodich, M., Gibson, M. C. and Layden, M. J. (2023). Environmental and molecular regulation of asexual reproduction in the sea anemone Nematostella vectensis. 2023.01.27.525773.

Ambrosone, A., Marchesano, V., Mazzarella, V. and Tortiglione, C. (2014). Nanotoxicology using the sea anemone *Nematostella vectensis*: from developmental toxicity to genotoxicology. Nanotoxicology 8, 508–520.

Amiel, A. R., Johnston, H. T., Nedoncelle, K., Warner, J. F., Ferreira, S. and Röttinger, E. (2015). Characterization of morphological and cellular events underlying oral regeneration in the sea anemone, *Nematostella vectensis*. International Journal of Molecular Sciences 16, 28449–28471.

Amiel, A. R., Foucher, K., Ferreira, S. and Röttinger, E. (2019). Synergic coordination of stem cells is required to induce a regenerative response in anthozoan cnidarians. bioRxiv 2019.12.31.891804.

Amiel, A. R., Michel, V., Carvalho, J. E., Shkreli, M., Petit, C. and Röttinger, E. (2021). The sea anemone *Nematostella vectensis*, an emerging model for biomedical research: Mechano-sensitivity, extreme regeneration and longevity. Med Sci (Paris) 37, 167–177.

Angilletta, M. J., Jr., Steury, T. D. and Sears, M. W. (2004). Temperature, growth rate, and body size in ectotherms: Fitting pieces of a life-history puzzle. Integrative and Comparative Biology 44, 498–509.

Arboleda, E., Hartenstein, V., Martinez, P., Reichert, H., Sen, S., Sprecher, S. and Bailly, X. (2018). An emerging system to study photosymbiosis, brain regeneration, chronobiology, and behavior: The marine Acoel *Symsagittifera roscoffensis*. BioEssays 40, 1800107.

Babonis, L. S., Enjolras, C., Reft, A. J., Foster, B. M., Hugosson, F., Ryan, J. F., Daly, M. and Martindale, M. Q. (2021). Knockout of a single Sox gene resurrects an ancestral cell type in the sea anemone Nematostella vectensis. 2021.09.30.462561.

Baldassarre, L., Ying, H., Reitzel, A. M., Franzenburg, S. and Fraune, S. (2022). Microbiota mediated plasticity promotes thermal adaptation in the sea anemone *Nematostella vectensis*. Nat Commun 13, 3804.

Best, J., Adatto, I., Cockington, J., James, A. and Lawrence, C. (2010). A novel method for rearing first-feeding larval zebrafish: Polyculture with type L saltwater rotifers (*Brachionus plicatilis*). Zebrafish 7, 289–295.

Bossert, P. E., Dunn, M. P. and Thomsen, G. H. (2013). A staging system for the regeneration of a polyp from the aboral physa of the anthozoan Cnidarian *Nematostella vectensis*. Developmental Dynamics 242, 1320–1331.

Bradford, Y. M., Van Slyke, C. E., Ruzicka, L., Singer, A., Eagle, A., Fashena, D., Howe, D. G., Frazer, K., Martin, R., Paddock, H., et al. (2022). Zebrafish information network, the knowledgebase for *Danio rerio* research. Genetics 220, iyac016.

Brix, K. V., Cardwell, R. D. and Adams, W. J. (2003). Chronic toxicity of arsenic to the Great Salt Lake brine shrimp, *Artemia franciscana*. Ecotoxicology and Environmental Safety 54, 169–175.

Busengdal, H. and Rentzsch, F. (2017). Unipotent progenitors contribute to the generation of sensory cell types in the nervous system of the cnidarian *Nematostella vectensis*. Developmental Biology 431, 59–68.

Carvalho, J. E., Lahaye, F. and Schubert, M. (2017). Keeping amphioxus in the laboratory: an update on available husbandry methods. Int. J. Dev. Biol. 61, 773–783.

Chames, P., Epinat, J.-C., Guillier, S., Patin, A., Lacroix, E. and Pâques, F. (2005). *In vivo* selection of engineered homing endonucleases using double-strand break induced homologous recombination. Nucleic Acids Research 33, e178.

Chen, C.-Y., McKinney, S. A., Ellington, L. R. and Gibson, M. C. (2020). Hedgehog signaling is required for endomesodermal patterning and germ cell development in the sea anemone *Nematostella vectensis*. eLife 9, e54573.

Chiruvella, K. K., Liang, Z. and Wilson, T. E. (2013). Repair of double-strand breaks by end joining. Cold Spring Harb Perspect Biol 5, a012757.

Chourrout, D., Delsuc, F., Chourrout, P., Edvardsen, R. B., Rentzsch, F., Renfer, E., Jensen, M. F., Zhu, B., de Jong, P., Steele, R. E., et al. (2006). Minimal ProtoHox cluster inferred from bilaterian and cnidarian Hox complements. Nature 442, 684–687.

Cook, C. E., Chenevert, J., Larsson, T. A., Arendt, D., Houliston, E. and Lénárt, P. (2016). Old knowledge and new technologies allow rapid development of model organisms. MBoC 27, 882–887.

Dabrowski, K. and Miller, M. (2018). Contested paradigm in raising Zebrafish (*Danio rerio*). Zebrafish 15, 295–309.

Davis, P., Zarowiecki, M., Arnaboldi, V., Becerra, A., Cain, S., Chan, J., Chen, W. J., Cho, J., da Veiga Beltrame, E., Diamantakis, S., et al. (2022). WormBase in 2022—data, processes, and tools for analyzing *Caenorhabditis elegans*. Genetics 220, iyac003.

Dhont, J., Dierckens, K., Støttrup, J., Van Stappen, G., Wille, M. and Sorgeloos, P. (2013). Rotifers, *Artemia* and copepods as live feeds for fish larvae in aquaculture. In Advances in Aquaculture Hatchery Technology (ed. Allan, G.) and Burnell, G.), pp. 157–202. Woodhead Publishing.

Doudna, J. A. and Charpentier, E. (2014). The new frontier of genome engineering with CRISPR-Cas9. Science 346, 1258096.

Ehrlich, W., Gahan, J. M., Rentzsch, F. and Kühn, F. J. P. (2022). TRPM2 causes sensitization to oxidative stress but attenuates high-temperature injury in the sea anemone *Nematostella vectensis*. Journal of Experimental Biology 225, jeb243717.

Elliott, B., Richardson, C., Winderbaum, J., Nickoloff, J. A. and Jasin, M. (1998). Gene conversion tracts from double-strand break repair in mammalian cells. Molecular and Cellular Biology 18, 93–101.

Elran, R., Raam, M., Kraus, R., Brekhman, V., Sher, N., Plaschkes, I., Chalifa-Caspi, V. and Lotan, T. (2014). Early and late response of *Nematostella vectensis* transcriptome to heavy metals. Molecular Ecology 23, 4722–4736.

Fei, J.-F., Lou, W. P.-K., Knapp, D., Murawala, P., Gerber, T., Taniguchi, Y., Nowoshilow, S., Khattak, S. and Tanaka, E. M. (2018). Application and optimization of CRISPR–Cas9-mediated genome engineering in axolotl (*Ambystoma mexicanum*). Nat Protoc 13, 2908– 2943.

Fritzenwanker, J. H. and Technau, U. (2002). Induction of gametogenesis in the basal cnidarian *Nematostella vectensis* (Anthozoa). Dev Genes Evol 212, 99–103.

Fritzenwanker, J. H., Genikhovich, G., Kraus, Y. and Technau, U. (2007). Early development and axis specification in the sea anemone *Nematostella vectensis*. Developmental Biology 310, 264–279.

Gahan, J. M., Kouzel, I. U., Jansen, K. O., Burkhardt, P. and Rentzsch, F. (2022). Histone demethylase Lsd1 is required for the differentiation of neural cells in *Nematostella vectensis*. Nat Commun 13, 465.

Genikhovich, G. and Technau, U. (2009). Induction of spawning in the starlet sea anemone *Nematostella vectensis*, in vitro fertilization of gametes, and dejellying of zygotes. Cold Spring Harb Protoc 2009, pdb.prot5281.

Gordon, T., Roth, L., Caicci, F., Manni, L. and Shenkar, N. (2020). Spawning induction, development and culturing of the solitary ascidian *Polycarpa mytiligera*, an emerging model for regeneration studies. Frontiers in Zoology 17, 19.

Gramates, L. S., Agapite, J., Attrill, H., Calvi, B. R., Crosby, M. A., dos Santos, G., Goodman, J. L., Goutte-Gattat, D., Jenkins, V. K., Kaufman, T., et al. (2022). FlyBase: a guided tour of highlighted features. Genetics 220, iyac035.

Hand, C. and Uhlinger, K. R. (1992). The culture, sexual and asexual reproduction, and growth of the sea anemone *Nematostella vectensis*. The Biological Bulletin 182, 169–176.

Hand, C. and Uhlinger, K. R. (1994). The unique, widely distributed, estuarine sea anemone, *Nematostella vectensis* Stephenson: A review, new facts, and questions. Estuaries 17, 501–508.

Havrilak, J. A., Faltine-Gonzalez, D., Wen, Y., Fodera, D., Simpson, A. C., Magie, C. R. and Layden, M. J. (2017). Characterization of *NvLWamide-like* neurons reveals stereotypy in *Nematostella* nerve net development. Developmental Biology 431, 336–346.

He, S., del Viso, F., Chen, C.-Y., Ikmi, A., Kroesen, A. E. and Gibson, M. C. (2018). An axial Hox code controls tissue segmentation and body patterning in *Nematostella vectensis*. Science 361, 1377–1380.

Henry, J. Q., Lesoway, M. P. and Perry, K. J. (2020). An automated aquatic rack system for rearing marine invertebrates. BMC Biology 18, 46.

Hill, E. M., Chen, C.-Y., del Viso, F., Ellington, L. R., He, S., Karabulut, A., Paulson, A. and Gibson, M. C. (2022). Manipulation of gene activity in the regenerative model Sea Anemone, *Nematostella vectensis*. In Whole-Body Regeneration: Methods and Protocols (ed. Blanchoud, S.) and Galliot, B.), pp. 437–465. New York, NY: Springer US.

Höijer, I., Emmanouilidou, A., Östlund, R., van Schendel, R., Bozorgpana, S., Tijsterman, M., Feuk, L., Gyllensten, U., den Hoed, M. and Ameur, A. (2022). CRISPR-Cas9 induces large structural variants at on-target and off-target sites in vivo that segregate across generations. Nat Commun 13, 627.

Holland, L. Z. and Li, G. (2021). Laboratory culture and mutagenesis of amphioxus (*Branchiostoma floridae*). In Developmental Biology of the Sea Urchin and Other Marine Invertebrates: Methods and Protocols (ed. Carroll, D. J.) and Stricker, S. A.), pp. 1–29. New York, NY: Springer US.

Ikmi, A. and Gibson, M. C. (2010). Identification and *in vivo* characterization of NvFP-7R, a developmentally regulated red fluorescent protein of *Nematostella vectensis*. PLOS ONE 5, e11807.

Ikmi, A., Delventhal, K. M., Gibson, M. C. and McKinney, S. A. (2014). TALEN and CRISPR/Cas9-mediated genome editing in the early-branching metazoan *Nematostella vectensis*. Nature Communications 5, 5486.

Ikmi, A., Steenbergen, P. J., Anzo, M., McMullen, M. R., Stokkermans, A., Ellington, L. R. and Gibson, M. C. (2020). Feeding-dependent tentacle development in the sea anemone *Nematostella vectensis*. Nat Commun 11, 4399.

Ivankovic, M., Haneckova, R., Thommen, A., Grohme, M. A., Vila-Farré, M., Werner, S. and Rink, J. C. (2019). Model systems for regeneration: planarians. Development 146, dev167684.

Johnston, H., Warner, J. F., Amiel, A. R., Nedoncelle, K., Carvalho, J. E. and Röttinger, E. (2021). Whole body regeneration deploys a rewired embryonic gene regulatory network logic. 658930.

Kassmer, S. H., Nourizadeh, S. and De Tomaso, A. W. (2019). Cellular and molecular mechanisms of regeneration in colonial and solitary Ascidians. Developmental Biology 448, 271–278.

Klein, S., Frazier, V., Readdean, T., Lucas, E., Diaz-Jimenez, E. P., Sogin, M., Ruff, E. S. and Echeverri, K. (2021). Common environmental pollutants negatively affect development and regeneration in the sea anemone *Nematostella vectensis* holobiont. Frontiers in Ecology and Evolution 9,.

Kraus, Y., Aman, A., Technau, U. and Genikhovich, G. (2016). Pre-bilaterian origin of the blastoporal axial organizer. Nat Commun 7, 11694.

Kusserow, A., Pang, K., Sturm, C., Hrouda, M., Lentfer, J., Schmidt, H. A., Technau, U., von Haeseler, A., Hobmayer, B., Martindale, M. Q., et al. (2005). Unexpected complexity of the *Wnt* gene family in a sea anemone. Nature 433, 156–160.

Lawrence, C. (2007). The husbandry of zebrafish (*Danio rerio*): A review. Aquaculture 269, 1–20.

Lawrence, C., Sanders, E. and Henry, E. (2012). Methods for culturing saltwater rotifers (*Brachionus plicatilis*) for rearing larval zebrafish. Zebrafish 9, 140–146.

Layden, M. J., Rentzsch, F. and Röttinger, E. (2016). The rise of the starlet sea anemone *Nematostella vectensis* as a model system to investigate development and regeneration. WIREs Dev Biol 5, 408–428.

Lebedeva, T., Boström, J., Mörsdorf, D., Niedermoser, I., Genikhovich, E., Adameyko, I. and Genikhovich, G. (2022). β-catenin-dependent endomesoderm specification appears to be a Bilateria-specific co-option. 2022.10.15.512282.

Lebouvier, M., Miramón-Puértolas, P. and Steinmetz, P. R. H. (2022). Evolutionary conserved aspects of animal nutrient uptake and transport in sea anemone vitellogenesis. 2022.01.24.477498.

Lechable, M., Jan, A., Duchene, A., Uveira, J., Weissbourd, B., Gissat, L., Collet, S., Gilletta, L., Chevalier, S., Leclère, L., et al. (2020). An improved whole life cycle culture protocol for the hydrozoan genetic model *Clytia hemisphaerica*. Biology Open 9, bio051268.

Lee, P. N., Kumburegama, S., Marlow, H. Q., Martindale, M. Q. and Wikramanayake, A. H. (2007). Asymmetric developmental potential along the animal–vegetal axis in the anthozoan cnidarian, *Nematostella vectensis*, is mediated by Dishevelled. Developmental Biology 310, 169–186.

Lieber, M. R. (2010). The mechanism of double-strand DNA break repair by the nonhomologous DNA end-joining pathway. Annual Review of Biochemistry 79, 181–211.

Lieschke, G. J. and Currie, P. D. (2007). Animal models of human disease: zebrafish swim into view. Nat Rev Genet 8, 353–367.

Mehta, A. S. and Singh, A. (2019). Insights into regeneration tool box: An animal model approach. Developmental Biology 453, 111–129.

Miyaoka, Y., Berman, J. R., Cooper, S. B., Mayerl, S. J., Chan, A. H., Zhang, B., Karlin-Neumann, G. A. and Conklin, B. R. (2016). Systematic quantification of HDR and NHEJ reveals effects of locus, nuclease, and cell type on genome-editing. Sci Rep 6, 23549.

Momose, T. and Concordet, J.-P. (2016). Diving into marine genomics with CRISPR/Cas9 systems. Marine Genomics 30, 55–65.

Murabe, N., Okumura, E., Chiba, K., Hosoda, E., Ikegami, S. and Kishimoto, T. (2021). The starfish *Asterina pectinifera*: Collection and maintenance of adults and rearing and metamorphosis of larvae. In Developmental Biology of the Sea Urchin and Other Marine Invertebrates: Methods and Protocols (ed. Carroll, D. J.) and Stricker, S. A.), pp. 49–68. New York, NY: Springer US.

Nakanishi, N. and Martindale, M. Q. (2018). CRISPR knockouts reveal an endogenous role for ancient neuropeptides in regulating developmental timing in a sea anemone. eLife 7, e39742.

Nakanishi, N., Renfer, E., Technau, U. and Rentzsch, F. (2012). Nervous systems of the sea anemone *Nematostella vectensis* are generated by ectoderm and endoderm and shaped by distinct mechanisms. Development 139, 347–357.

Ormestad, M., Martindale, M. Q. and Röttinger, E. (2011). A comparative gene expression database for invertebrates. EvoDevo 2, 17.

Paix, A., Basu, S., Steenbergen, P., Singh, R., Prevedel, R. and Ikmi, A. (2023). Endogenous tagging of multiple cellular components in the sea anemone *Nematostella vectensis*. Proceedings of the National Academy of Sciences 120, e2215958120.

Pan, Y.-J., Dahms, H.-U., Hwang, J.-S. and Souissi, S. (2022). Recent trends in live feeds for marine larviculture: A mini review. Frontiers in Marine Science 9,.

Passamaneck, Y. J. and Martindale, M. Q. (2012). Cell proliferation is necessary for the regeneration of oral structures in the anthozoan cnidarian *Nematostella vectensis*. BMC Developmental Biology 12, 34.

Reitzel, A. M., Burton, P. M., Krone, C. and Finnerty, J. R. (2007). Comparison of developmental trajectories in the starlet sea anemone *Nematostella vectensis*: embryogenesis, regeneration, and two forms of asexual fission. Invertebrate Biology 126, 99–112.

Reitzel, A. M., Sullivan, J. C., Traylor-knowles, N. and Finnerty, J. R. (2008). Genomic survey of candidate stress-response genes in the estuarine anemone *Nematostella vectensis*. The Biological Bulletin 214, 233–254.

Renfer, E., Amon-Hassenzahl, A., Steinmetz, P. R. H. and Technau, U. (2010). A muscle-specific transgenic reporter line of the sea anemone, *Nematostella vectensis*. Proceedings of the National Academy of Sciences 107, 104–108.

Reuven, S., Rinsky, M., Brekhman, V., Malik, A., Levy, O. and Lotan, T. (2021). Cellular pathways during spawning induction in the starlet sea anemone *Nematostella vectensis*. Sci Rep 11, 15451.

Richards, G. S. and Rentzsch, F. (2014). Transgenic analysis of a *SoxB* gene reveals neural progenitor cells in the cnidarian *Nematostella vectensis*. Development 141, 4681–4689.

Röttinger, E. (2021). *Nematostella vectensis*, an emerging model for deciphering the molecular and cellular mechanisms underlying whole-body regeneration. Cells 10, 2692.

Röttinger, E., Dahlin, P. and Martindale, M. Q. (2012). A framework for the establishment of a cnidarian gene regulatory network for “endomesoderm” specification: The inputs of ß-Catenin/TCF signaling. PLOS Genetics 8, e1003164.

Roux, N., Salis, P., Lee, S.-H., Besseau, L. and Laudet, V. (2020). Anemonefish, a model for Eco-Evo-Devo. EvoDevo 11, 20.

Schneider, C. A., Rasband, W. S. and Eliceiri, K. W. (2012). NIH Image to ImageJ: 25 years of image analysis. Nat Meth 9, 671–675.

Seleit, A., Aulehla, A. and Paix, A. (2021). Endogenous protein tagging in medaka using a simplified CRISPR/Cas9 knock-in approach. eLife 10, e75050.

Servetnick, M. D., Steinworth, B., Babonis, L. S., Simmons, D., Salinas-Saavedra, M. and Martindale, M. Q. (2017). Cas9-mediated excision of *Nematostella brachyury* disrupts endoderm development, pharynx formation and oral-aboral patterning. Development 144, 2951–2960.

Silva, G., Poirot, L., Galetto, R., Smith, J., Montoya, G., Duchateau, P. and Paques, F. (2011). Meganucleases and other tools for targeted genome engineering: Perspectives and challenges for gene therapy. Current Gene Therapy 11, 11–27.

Soto-Àngel, J. J., Nordmann, E.-L., Sturm, D., Sachkova, M., Pang, K. and Burkhardt, P. (2022). Stable laboratory culture system for the ctenophore Mnemiopsis leidyi.

Srivastava, M. (2022). Studying development, regeneration, stem cells, and more in the acoel *Hofstenia miamia*. In Current Topics in Developmental Biology (ed. Goldstein, B.) and Srivastava, M.), pp. 153–172. Academic Press.

Stefanik, D. J., Friedman, L. E. and Finnerty, J. R. (2013). Collecting, rearing, spawning and inducing regeneration of the starlet sea anemone, *Nematostella vectensis*. Nat Protoc 8, 916–923.

Steger, J., Cole, A. G., Denner, A., Lebedeva, T., Genikhovich, G., Ries, A., Reischl, R., Taudes, E., Lassnig, M. and Technau, U. (2022). Single-cell transcriptomics identifies conserved regulators of neuroglandular lineages. Cell Reports 40, 111370.

Steinmetz, P. R. H., Aman, A., Kraus, J. E. M. and Technau, U. (2017). Gut-like ectodermal tissue in a sea anemone challenges germ layer homology. Nature Ecology & Evolution 1, 1535–1542.

Sunagar, K., Columbus-Shenkar, Y. Y., Fridrich, A., Gutkovich, N., Aharoni, R. and Moran, Y. (2018). Cell type-specific expression profiling unravels the development and evolution of stinging cells in sea anemone. BMC Biology 16, 108.

Tkavc, R., Ausec, L., Oren, A. and Gunde-Cimerman, N. (2011). Bacteria associated with *Artemia* spp. along the salinity gradient of the solar salterns at Eilat (Israel). FEMS Microbiology Ecology 77, 310–321.

Tournière, O., Dolan, D., Richards, G. S., Sunagar, K., Columbus-Shenkar, Y. Y., Moran, Y. and Rentzsch, F. (2020). *NvPOU4/Brain3* functions as a terminal selector gene in the nervous system of the Cnidarian *Nematostella vectensis*. Cell Reports 30, 4473–4489.e5.

Valverde, E. J., Labella, A. M., Borrego, J. J. and Castro, D. (2019). *Artemia* spp., a susceptible host and vector for lymphocystis disease virus. Viruses 11, 506.

Varshney, G. K., Carrington, B., Pei, W., Bishop, K., Chen, Z., Fan, C., Xu, L., Jones, M., LaFave, M. C., Ledin, J., et al. (2016). A high-throughput functional genomics workflow based on CRISPR/Cas9-mediated targeted mutagenesis in zebrafish. Nat Protoc 11, 2357– 2375.

Warner, J. F., Guerlais, V., Amiel, A. R., Johnston, H., Nedoncelle, K. and Röttinger, E. (2018). NvERTx: a gene expression database to compare embryogenesis and regeneration in the sea anemone *Nematostella vectensis*. Development 145,.

Wijesena, N., Simmons, D. K. and Martindale, M. Q. (2017). Antagonistic BMP–cWNT signaling in the cnidarian *Nematostella vectensis* reveals insight into the evolution of mesoderm. Proceedings of the National Academy of Sciences 114, E5608–E5615.

Wikramanayake, A. H., Hong, M., Lee, P. N., Pang, K., Byrum, C. A., Bince, J. M., Xu, R. and Martindale, M. Q. (2003). An ancient role for nuclear β-catenin in the evolution of axial polarity and germ layer segregation. Nature 426, 446–450.

Williams, R. B. (1983). Starlet sea anemone. IUCN Invertebrate Red Data Book 43–46.

Yoshimi, K., Oka, Y., Miyasaka, Y., Kotani, Y., Yasumura, M., Uno, Y., Hattori, K., Tanigawa, A., Sato, M., Oya, M., et al. (2021). *Combi*-CRISPR: combination of NHEJ and HDR provides efficient and precise plasmid-based knock-ins in mice and rats. Hum Genet 140, 277–287.

Zang, H. and Nakanishi, N. (2020). Expression analysis of cnidarian-specific neuropeptides in a sea anemone unveils an apical-organ-associated nerve net that disintegrates at metamorphosis. Frontiers in Endocrinology 11,.

Zhang, J.-P., Li, X.-L., Li, G.-H., Chen, W., Arakaki, C., Botimer, G. D., Baylink, D., Zhang, L., Wen, W., Fu, Y.-W., et al. (2017). Efficient precise knockin with a double cut HDR donor after CRISPR/Cas9-mediated double-stranded DNA cleavage. Genome Biology 18, 35.

Zimmermann, B., Robb, S. M. C., Genikhovich, G., Fropf, W. J., Weilguny, L., He, S., Chen, S., Lovegrove-Walsh, J., Hill, E. M., Chen, C.-Y., et al. (2022). Sea anemone genomes reveal ancestral metazoan chromosomal macrosynteny. 2020.10.30.359448.

